# Inhibition of the 60S ribosome biogenesis GTPase LSG1 causes endoplasmic reticular disruption and cellular senescence

**DOI:** 10.1101/463851

**Authors:** Asimina Pantazi, Andrea Quintanilla, Priya Hari, Nuria Tarrats, Eleftheria Parasyraki, Flora Lucy Dix, Jaiyogesh Patel, Tamir Chandra, Juan Carlos Acosta, Andrew John Finch

## Abstract

Cellular senescence is triggered by diverse stimuli and is characterised by long-term growth arrest and secretion of cytokines and chemokines (termed the SASP - senescence-associated secretory phenotype). Senescence can be organismally beneficial as it can prevent the propagation of damaged or mutated clones and stimulate their clearance by immune cells. However, it has recently become clear that senescence also contributes to the pathophysiology of aging through the accumulation of damaged cells within tissues. Here we describe that inhibition of the reaction catalysed by LSG1, a GTPase involved in the biogenesis of the 60S ribosomal subunit, leads to a robust induction of cellular senescence. Perhaps surprisingly, this was not due to ribosome depletion or translational insufficiency, but rather through perturbation of endoplasmic reticulum (ER) homeostasis and a dramatic upregulation of the cholesterol biosynthesis pathway. This cholesterol/ER signature is shared with several other forms of senescence and contributes to the cell cycle arrest in oncogene-induced senescence (OIS). Furthermore, targetting of LSG1 resulted in amplification of the cholesterol/ER signature and restoration of a robust cellular senescence response in transformed cells, suggesting potential therapeutic uses of LSG1 inhibition.

## Introduction

Mammalian ribosomes are nucleoprotein complexes comprised of a large (60S) subunit and a small (40S) subunit that carry out the fundamental process of translation. The mature ribosome contains four ribosomal RNAs (rRNAs) and almost 80 proteins and the complex process of ribosome biogenesis involves over 200 trans-acting factors (reviewed in (Kressler, Hurt, & Baßler, 2017)). Transcription of rRNA precursors from tandem repeats of ribosomal DNA (rDNA) initiates ribosome biogenesis and a complex sequence of events including sequential splicing and recruitment of rRNA-associated proteins ensues. Mutations in genes that encode core ribosomal proteins or factors involved in ribosome biogenesis give rise to diseases that are collectively termed ribosomopathies. Examples of these inherited disorders include Treacher Collins Syndrome, Diamond-Blackfan anaemia and Shwachman Diamond Syndrome (reviewed in (Danilova & Gazda, 2015)). The acquired myelodysplastic syndrome 5q-, characterised by a deletion of a region of chromosome 5q, is also considered a ribosomopathy due to the presence of the *RPS14* gene in the deleted region and the phenotypic recapitulation of much of the disease phenotype upon deletion of *RPS14* alone (Barlow et al., 2010; Ebert et al., 2008). Given the requirement for ribosome biogenesis in cellular growth and proliferation, the causative mutation in these diseases is clearly detrimental to the cell. However, the pathology that arises in these ribosomopathies is, in many cases, caused by activation of the p53 pathway in response to the primary lesions ((Barkic et al., 2009; Barlow et al., 2010; Jones et al., 2008). The exact nature of the stresses that activate the p53 pathway in the ribosomopathies remains undefined.

Regulation of ribosome biogenesis occurs primarily at the level of the transcriptional complexes that are recruited to the rDNA. The majority of rRNA is produced by RNA polymerase I-mediated transcription and this activity requires recruitment of TIF-1A (transcription initiation factor 1A), UBF (upstream binding factor) and SL1 (selectivity factor 1) to rDNA promoter regions. Both of these factors are regulated by phosphorylation and they thereby integrate signals from the MAP kinase and mTOR pathways (Hannan et al., 2003; Mayer, Zhao, Yuan, & Grummt, 2004; Zhao, Yuan, Frödin, & Grummt, 2003). In addition, UBF is activated through interaction with c-Myc (Poortinga et al., 2004) and inhibited by the Rb (Cavanaugh et al., 1995; Voit, Schäfer, & Grummt, 1997) and p53 (Budde & Grummt, 1999; Zhai & Comai, 2000) pathways. Accordingly, deregulation of ribosome biogenesis is commonly seen in cancer and the histochemical AgNOR test (for silver-binding <u>A</u>r<u>G</u>yrophilic <u>N</u>ucleolar <u>O</u>rganiser <u>R</u>egions) is used for staging and prognosis in many cases (Pich, Chiusa, & Margaria, 2000). The increased ribosome biogenesis observed in cancer has encouraged the idea that inhibition of ribosome biogenesis could represent a therapeutic strategy in cancer therapy. Indeed, a small molecule inhibitor of RNA polymerase I, CX-5461, has recently shown promise in this regard (Bywater et al., 2012; Drygin et al., 2011).

We identified the GTPases involved in the cytoplasmic maturation of the 60S ribosomal subunit as plausible targets for therapeutic intervention. These GTPases catalyse the release of two anti-association factors that are loaded onto the 60S particle in the nucleus and that are removed in the cytosol at the last stages of 60S maturation (Finch et al., 2011; Lo et al., 2010) (Fig 1a). EFL1 leads to eviction of the antiassociation factor eIF6 from the pre-60S in a reaction that requires the SBDS cofactor and GTP hydrolysis (Finch et al., 2011), whilst LSG1 catalyses the eviction of NMD3 in a reaction requiring RPL10, which stays associated with the ribosome (Hedges, West, & Johnson, 2005; Ma et al., 2017; Malyutin, Musalgaonkar, Patchett, Frank, & Johnson, 2017). Following removal of the two anti-association factors, the mature 60S subunit can then join the translating pool of ribosomes and the anti-association factors are returned to the nucleus to participate in subsequent rounds of 60S biogenesis.

**Fig 1.**
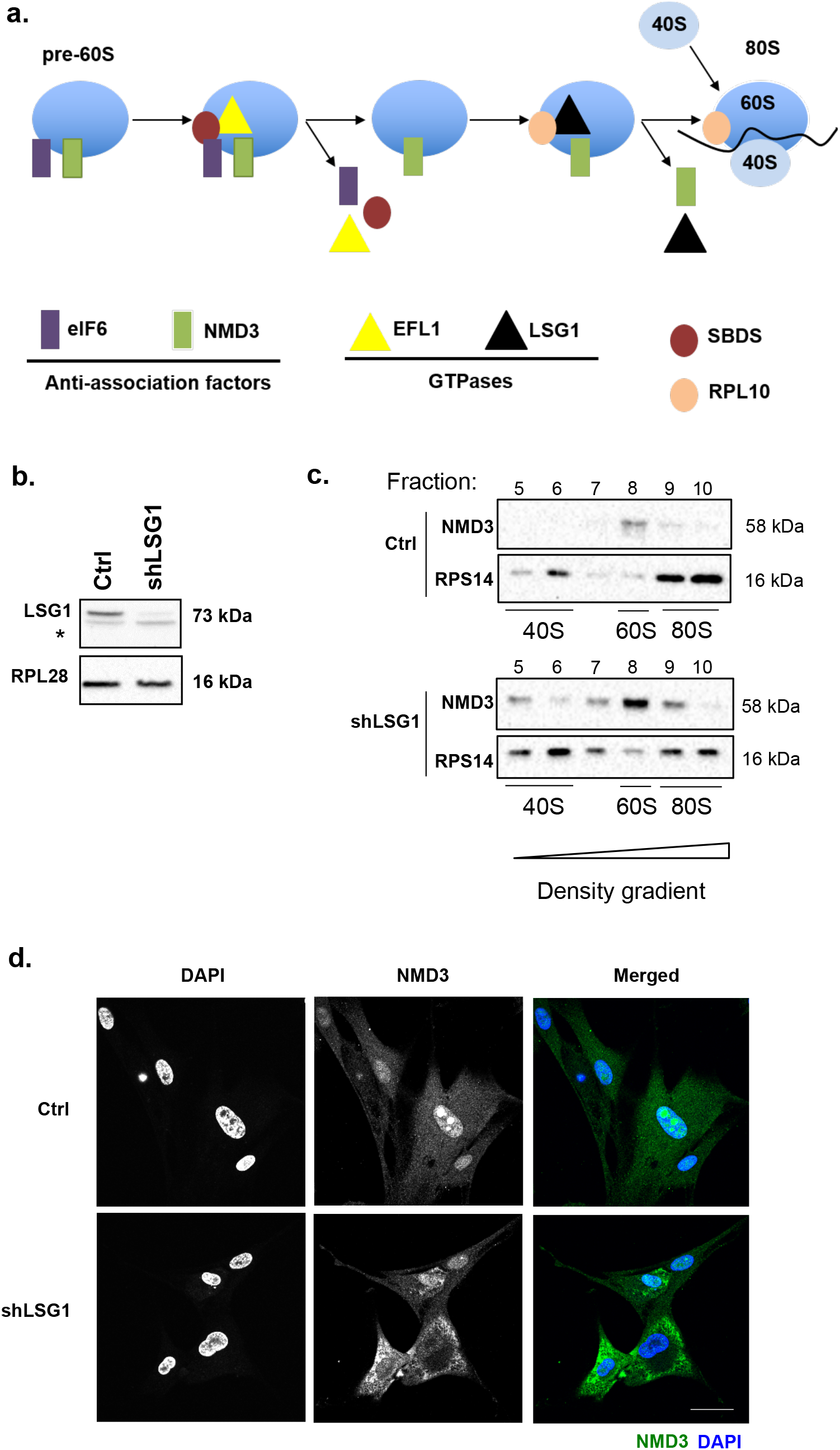
Knockdown of LSG1 inhibits NMD3 release from the ribosomal 60S subunit. a. Schematic of the late cytoplasmic reactions of 60S subunit maturation. The cytoplasmic pre-60S subunit carries the anti-association factors eIF6 and NMD3. Recruitment of the factor SBDS and the GTPase EFL1 leads to eviction of eIF6 in a reaction catalysed by hydrolysis of GTP. SBDS stimulates GTP hydrolysis by EFL1, which induces a rotation in the structure of SBDS, resulting in conformational changes and eIF6 release. RPL10 and the GTPase LSG1 then bind to the subunit leading to eviction of NMD3, again catalysed by GTP hydrolysis. RPL10 is retained on the 60S subunit and the mature 80S ribosome is formed. (Adapted from Hedges et al., 2005 and Finch et al., 2011)
b. Western blot analysis shows the knockdown of LSG1 in HEK 293 cells. The asterisk denotes a non-specific band. RPL28 was used as a reference protein.
c. Western blot analysis shows the levels of NMD3 and RPS14 across sequential fractions (5-10) collected from sucrose gradients in control and shLSG1 conditions. The NMD3 in fraction 8 corresponds to the localisation of 60S monomers. RPS14 indicates the localisation of the 40S and 80S components.
d. Immunostaining for NMD3 in MRC5 cells (control and shLSG1), followed by confocal microscopy, reveals accumulation of NMD3 in the cytoplasm following LSG1 knockdown. Scale bar: 50 μm

Here we report that knockdown of LSG1 and other components of the 60S maturation pathway promote a robust activation of cellular senescence. This senescence response is characterised by activation of the p53 and p16/Rb pathways and by a highly restricted SASP lacking the NF-κB-driven proinflammatory cytokines and chemokines. shLSG1 also promotes a striking upregulation of the cholesterol biosynthesis pathway and genes involved in endoplasmic reticulum (ER) organisation and this is accompanied by a disruption of the reticular morphology of the ER. Indeed, RPL10 and LSG1 have been shown to associate with ribosomes at the rough ER (Loftus, Nguyen, & Stanbridge, 1997; Reynaud et al., 2005) and our data suggest that loss of LSG1 significantly impacts upon ER homeostasis. Finally, we provide evidence that inhibition of 60S maturation can restore a robust senescence response in oncogene-transformed cells that have already bypassed oncogene-induced senescence.

## Results

### Knockdown of LSG1 inhibits NMD3 release from the ribosomal 60S subunit

The enzymes that catalyse the final cytoplasmic reactions in the maturation of the large (60S) ribosomal subunit (Fig 1a) represent possible targets for therapeutic inhibition. Accordingly, we chose RNAi rather than gene deletion as a strategy to mimic pharmacological inhibition because it can be efficient, yet not absolute. Since LSG1 catalyses the release of NMD3 from the cytoplasmic pre-60S particle, knockdown of LSG1 should result in failure to release NMD3 and thus to its cytosolic sequestration (Hedges et al., 2005; Ma et al., 2017; Malyutin et al., 2017). Infection of cells with lentiviral vector encoding a shRNA to LSG1 led to efficient knockdown of the protein, as assessed by western blot (Fig 1b) and, in turn, this led to an increase in association of NMD3 with the 60S subunit fraction as assessed by sucrose density gradient separation of ribosomal subunits (Fig1c). NMD3 is loaded onto pre-60S subunits in the nucleus (Gadal et al., 2001; Ho, Kallstrom, & Johnson, 2000) and immunofluorescent staining of control cells shows a nuclear/nucleolar staining pattern for NMD3. Knockdown of LSG1 led to relocalisation of NMD3 to the cytoplasm (Fig 1d), consistent with its retention on maturing cytoplasmic pre-60S particles due to loss of LSG1-mediated release. This relocalisation of Nmd3 from nucleus to cytoplasm is also observed in yeast lacking Lsg1 (Hedges et al., 2005) and is diagnostic of the defect in this maturation reaction.

### Impairment of 60S biogenesis triggers a robust cellular senescence response

We generated additional shRNAs to SBDS (the cofactor for EFL1 (Finch et al., 2011)) to target 60S maturation and assessed their knockdown by western blot: we obtained two shRNAs that were efficient for SBDS (Fig 2a). We introduced the shRNAs into primary human MRC5 fibroblasts through lentiviral transduction to impair 60S biogenesis and assessed their growth. Several days after viral infection, we noticed that impairment of 60S biogenesis led to a sparse culture and spreading of the cells with morphology that resembled cellular senescence. Analysis of BrdU incorporation using high content microscopy revealed that knockdown of LSG1 and SBDS led to a potent cell cycle arrest (Fig 2b, Sup Fig 1a) and this was accompanied by activation of acidic β-galactosidase activity and accumulation of p16 mRNA and protein (Fig 2c, Sup Figs 1a, 1b, 2a), indicating a senescence response. In addition to these core markers of cellular senescence we also observed increased staining for p53, p21 and the DNA damage response marker pST/Q (Sup Fig 1a, 2b). Furthermore, although the shSBDS(b) shRNA gave a less robust response with p53 and p21 immunofluorescence, p21 mRNA was induced consistent with activation of the p53 pathway (Sup Fig 2a,b). Senescence is characterized by ongoing, rather than transient, growth arrest and we confirmed the continuous nature of the shLSG1-induced growth defect through assessment of BrdU incorporation and p16, p53 and p21 immunoreactivity in a timecourse over 15 days (Fig 1d, Sup Fig 2c). To confirm the specificity for LSG1 in this process, we first generated and utilised a second shRNA to LSG1 (Sup Fig 3a) and again observed induction of acidic β-galactosidase activity (Sup Fig 3b) and reduction in BrdU staining (Sup Fig 3c). Next we used siRNA SMARTpools to EFL1 and LSG1 (Sup Fig 4a) and observed the expected reduction in BrdU incorporation (Sup Fig 4b) and induction of acidic β-galactosidase activity (Sup Fig 4c) and p16 immunoreactivity (Sup 4d). Deconvolution of the LSG1 siRNA pools revealed two independent siRNAs that knocked down LSG1 (Sup 5a) and reduced BrdU incorporation and induction of p16 and p53 (Sup Fig 5b). Taken together, these results demonstrate that inhibition of 60S ribosomal subunit maturation triggers a robust cellular senescence response.

**Fig 2.**
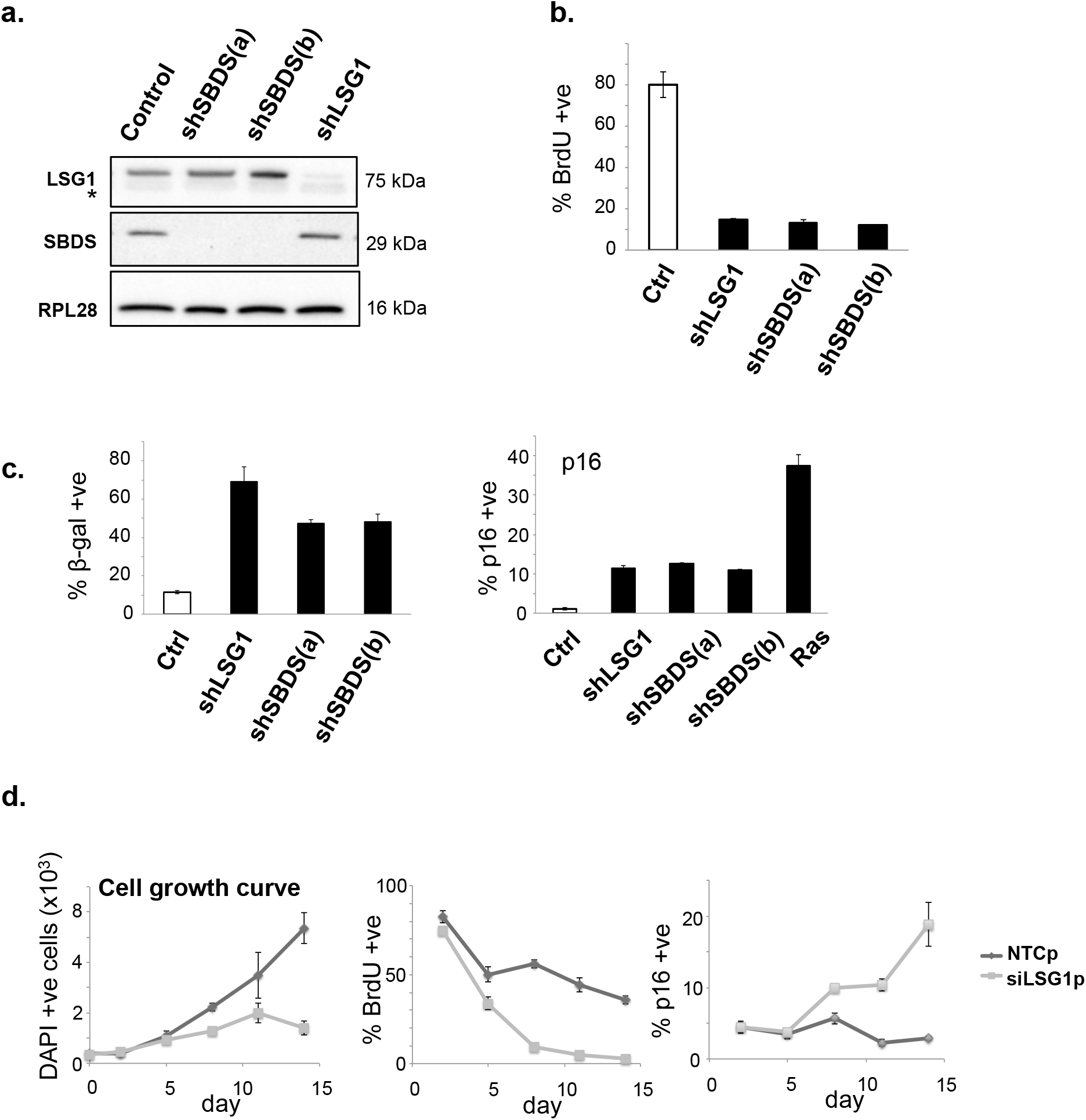
Knockdown of LSG1 and SBDS induces senescence. a. Western blot showing the efficiency of LSG1 and SBDS knockdown in MRC5 cells induced by the hairpins shLSG1, shSBDS(a) and shSBDS(b). The asterisk denotes a non-specific band in the LSG1 blot. RPL28 was used as a reference protein.
b. High content imaging analysis of BrdU incorporation and immunostaining in MRC5 cells with LSG1 and SBDS downregulation, 7 days post-infection. The cells were treated with 50mM BrdU for 16 hours.
c. The Senescence-Associated β-galactosidase assay was performed 7 days post-infection. Images were taken using phase contrast microscopy and the number of cells that were positive for the blue precipitate was counted. The bar chart on the right shows high content imaging analysis of p16 immunostaining. Ras-transduced cells were used as a positive control for p16 induction.
d. Timecourse experiment (timepoints: d0, d2, d5, d8, d11, d14) using a siRNA SmartPool for LSG1 (siLSG1p). Cell growth (DAPI stain), BrdU incorporation and p16 expression were monitored throughout the timecourse using high content microscopy. Error bars show standard deviation of 3 biological replicates.

### The senescence response to inhibition of 60S maturation is p53-dependent in primary cells

The two main pathways that implement most aspects of replicative and oncogene-induced senescence responses are the p16/retinoblastoma (RB) and p53 pathways (Salama, Sadaie, Hoare, & Narita, 2014). As described above, we observed that both pathways were activated by knockdown of LSG1 and we set out to determine which of these pathways was required for induction of senescence under this condition. The viral oncoproteins E6 and E7 from the human papilloma virus inhibit the p53 and Rb pathways, respectively, and are well-established tools for the determination of function of these pathways. We infected primary human fibroblasts with retroviral vectors expressing E6, E7 or an E6-E7 fusion protein (Acosta et al., 2008) and then with lentiviral shRNA to LSG1. Both viral constructs were functional since expression of E6 abrogated p53 expression whilst E7 expression enhanced p53 levels as previously described (Demers, Halbert, & Galloway, 1994)(Fig 3a). Loss of p53 function leads to bypass of replicative and oncogene-induced senescence (Bond, Wyllie, & Wynford-Thomas, 1994; Serrano, Lin, McCurrach, Beach, & Lowe, 1997) and, similarly, E6 expression led to continued BrdU incorporation upon LSG1 knockdown (Fig 3b). Expression of E7, on the other hand, did not rescue the inhibition of cell cycle elicited by LSG1 knockdown. We confirmed the p53-dependence of the senescence response using a C-terminal, dominant-negative fragment of p53, which leads to stabilization of the endogenous p53 protein through inhibition of its function (Fig 3c). Once again, inhibition of the p53 pathway led to bypass of shLSG1-induced proliferative arrest (Fig 3d). Finally, we used shRNA to p53 to follow the growth characteristics of cell lines transduced with shRNA to LSG1 (Fig 3e). p53 knockdown resulted in greatly accelerated growth rates in vector control cells and the knockdown of LSG1 failed to inhibit growth in these cells (Fig 3f).

**Fig 3.**
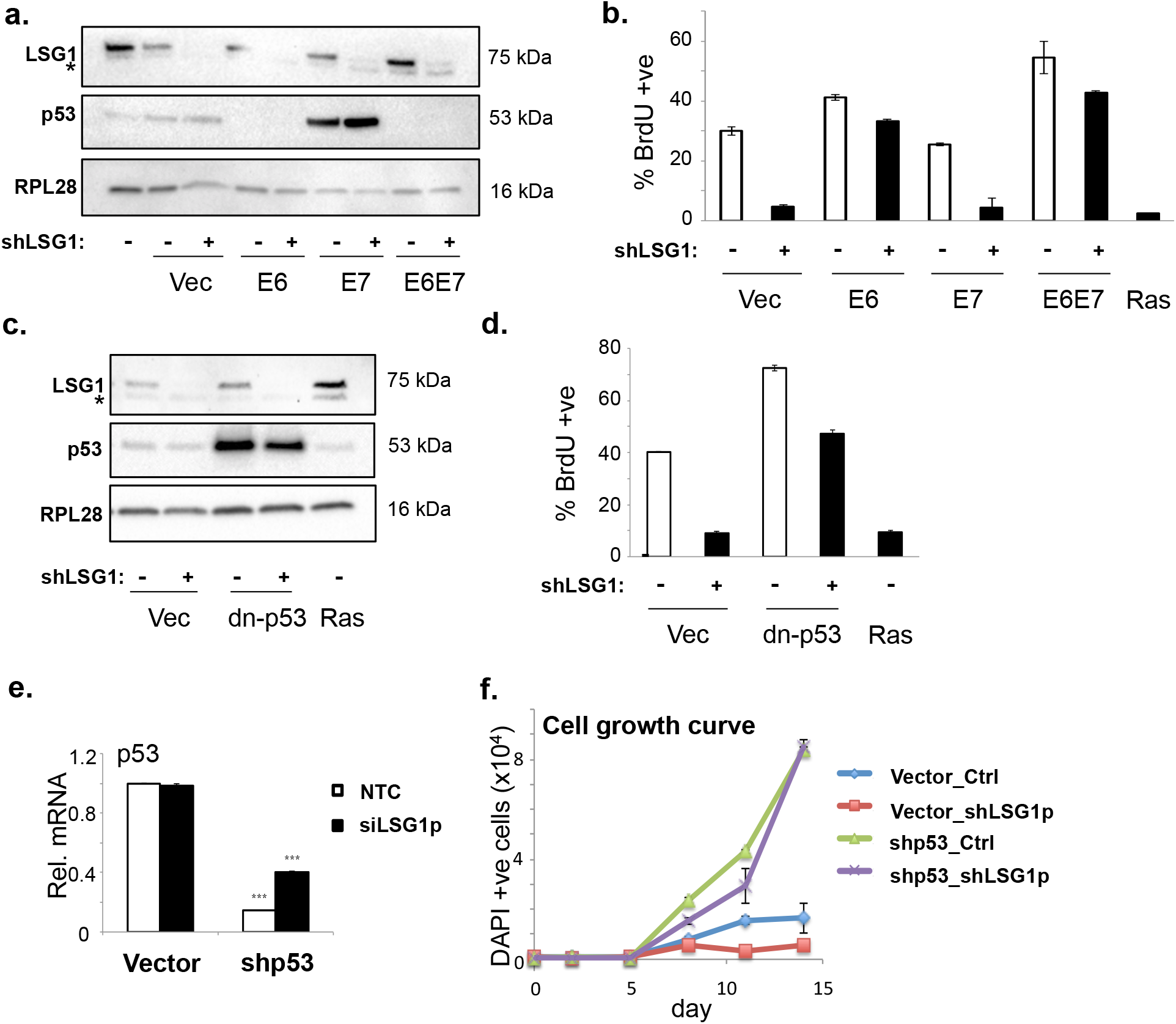
The senescence response induced by LSG1 knockdown is p53-dependent. a. Western blots for LSG1, p53 and RPL28 in MRC5 cells transduced with shLSG1 and/or HPV E6, E7 or E6E7 (the asterisk denotes a non-specific band).
b. BrdU incorporation was measured by high content imaging in cells transduced as in (a) above. K-RAS^V12^-transduced cells were used as a positive control for growth arrest.
c. Western blots for LSG1, p53 and RPL28 in MRC5 cells transduced with shLSG1 and/or a dominant negative p53 construct (dn-p53) (the asterisk denotes a non-specific band). Ras retroviral overexpression is included as a positive control.
d. BrdU incorporation was measured by high content imaging in cells transduced as in (c) above.
e. qPCR analysis of MRC5 cells transduced with shLSG1 and/or shp53 for the quantification of p53 transcript levels.
f. Timecourse experiment for the study of the growth levels of the above (e) cells, using high content imaging to measure DAPI stain. Timepoints: d0, d2, d5, d8, d11, d14. Error bars show standard deviation of 3 biological replicates.

### Inhibition of 60S maturation induces a senescent transcriptional response

The transcriptional responses to several triggers of senescence have recently been reported (Juan CAcosta et al., 2008; Juan Carlos Acosta et al., 2013; Hoare et al., 2016; Muñoz-Espín et al., 2013). Therefore, in order to gain mechanistic insight into the molecular cause of the senescence elicited by inhibition of 60S maturation, we performed transcriptomic analysis of shLSG1 cells. In particular, we sought to investigate molecular signatures that were shared with other forms of senescence. Senescence was induced in primary human MRC5 fibroblasts through transduction of shLSG1 and through overexpression of K-RasV12 as a positive control for oncogene-induced senescence. Extraction of RNA was followed by AmpliSeq library preparation and IonTorrent sequencing of amplicons. Global gene expression clustering revealed clear differences between the two senescent states (Fig 4a) and we therefore assessed similarity between the senescence caused by LSG1 knockdown and previously reported triggers of senescence. We generated gene sets from several systems in which senescence was induced, including OIS (Juan Carlos Acosta et al., 2013; Pawlikowski et al., 2013), replicative senescence (Pazolli et al., 2009), paracrine senescence (Juan Carlos Acosta et al., 2013), drug-induced senescence (Jing et al., 2011) and pancreatic intraepithelial neoplasia (Ling et al., 2012). These gene sets were used to measure enrichment in the shLSG1 dataset and in almost all cases showed enrichment with a false discovery rate-adjusted (FDR) Q-value of 0.01 or below (Fig 4b). The two exceptions that did not show statistically significant enrichment were developmental senescence (Muñoz-Espín et al., 2013) and DNA damage-induced senescence in hepatic stellate cells (Krizhanovsky et al., 2008) (Sup. Fig 6a). Thus the transcriptional response to LSG1 knockdown contains a strong senescent signature that is shared with multiple forms of senescence that arise *in vitro* and *in vivo*.

**Fig 4.**
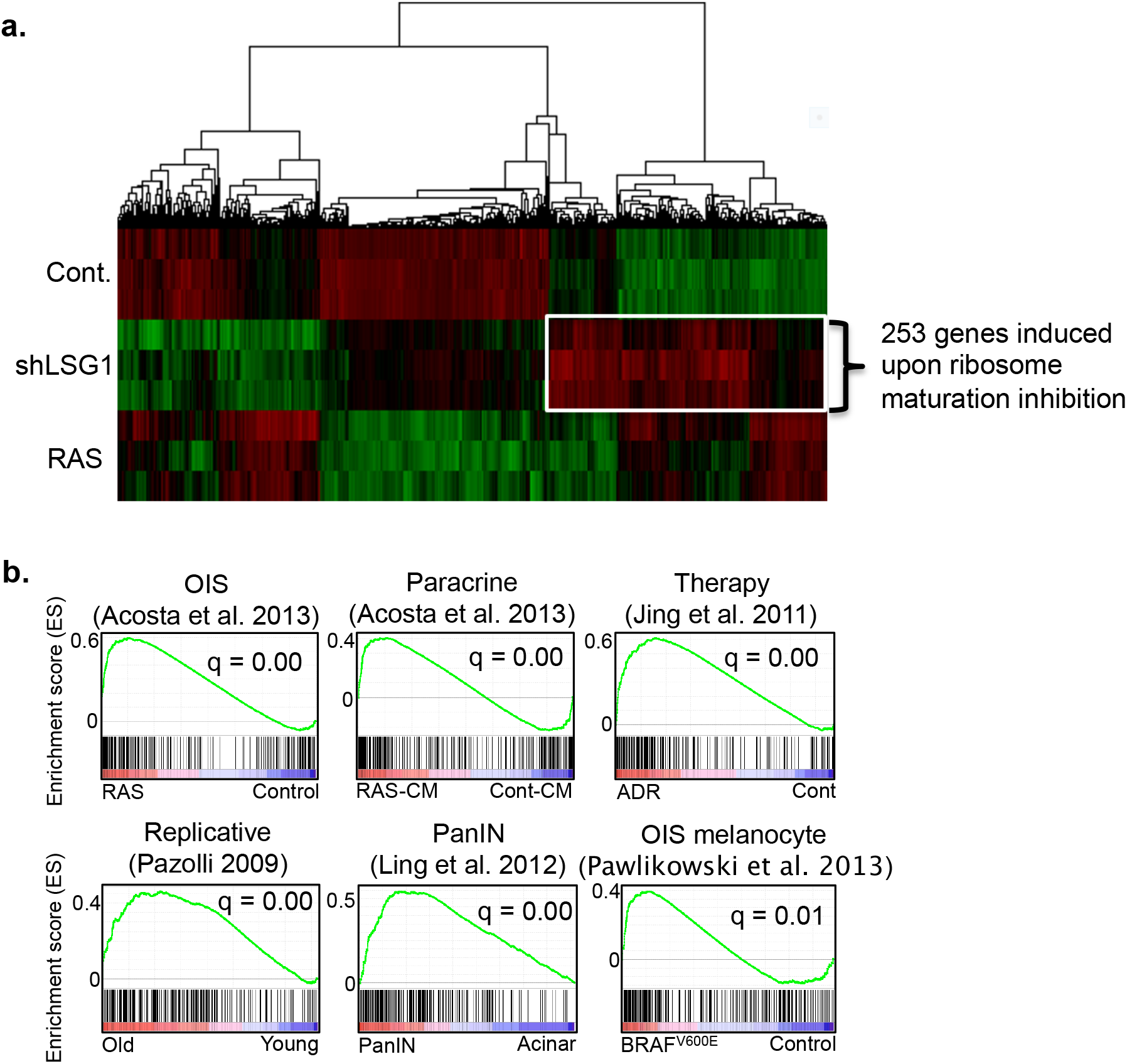
A signature of genes induced by LSG1 knockdown is common with other senescence responses. a. Hierarchical clustering of mRNA profiles from cells transduced with K-RAS^V12^, shLSG1 and vector control (Cont.) in MRC5 cells showing genes changing significantly (Adj.p<0.01) between shLSG1 and control. A signature of 253 genes induced by shLSG1 is highlighted. Data represents 3 experimental replicates. This analysis was performed using Cluster 3 and TreeView software.
b. GSEA plots showing that a signature of 253 genes derived from MRC5 cells undergoing shLSG1-induced senescence (described in a.) is found significantly enriched in multiple forms of senescence. (q represents false discovery rate (FDR)).

### 60S maturation inhibition induces production of a restricted SASP

As expected, we observed a marked antiproliferative signature characterised by upregulation of CDK inhibitors and downregulation of *E2F1*, cyclins and cyclin-dependent kinases (Fig 5a). One of the hallmarks of senescent cells is the release of a cocktail of pro-inflammatory cytokines and chemokines, collectively termed the senescence associated secretory phenotype (SASP). We analysed our transcriptomic data in more detail for genes previously identified as SASP-related or generally involved in inflammation (Juan Carlos Acosta et al., 2013). This analysis revealed a lack of most of the canonical SASP factors involved in OIS and revealed three distinct gene clusters (Fig 5b) – including one that was upregulated upon LSG1 knockdown but only weakly (or not at all) with OIS (Fig 5c, cluster 2) and one specific for OIS (Fig 5c, cluster 3). The shLSG1-specific cluster (cluster 2) included *TGFβ2* and *TGFβR1* as well as the other TGFβ family receptors *ACVR1* and *ACVR2a* and the TGFβ target genes *SERPINE1* and *IGFBP7* (Fig 5c). This was supported by gene set enrichment analysis which indicated significant enrichment of genes associated with the TGFβ signalling pathway (Fig 5d) and qRT-PCR analyses that verified upregulation of *SERPINE1*, *TGFB2* and *IGFBP7* (Sup Fig 6b). The OIS-specific transcriptome cluster included strongly pro-inflammatory cytokines and chemokines (Fig 5c, cluster 3, region A), indicative of the strong NF-κB-driven SASP program in OIS. Gene set enrichment analysis between the shLSG1 transcriptome and the OIS NF-κB programme showed no significant induction of these genes upon knockdown of LSG1 and this lack of key NF-κB-driven SASP components was confirmed at the mRNA (Fig 5e) and protein (Fig 5f) levels. In OIS, the SASP participates in autocrine/paracrine loops to reinforce the senescent phenotype (Juan CAcosta et al., 2008; Juan Carlos Acosta et al., 2013). We therefore targeted two of the components from our restricted SASP, namely *ACVR1B* and *TGFBR1*, to assess their contribution to the senescence response. shRNA to both factors resulted in reduction of their gene expression, although knockdown of *ACVR1B* was partial (Sup Fig 6c). Since TGFβ signaling induces senescence primarily through p15 expression (Hannon & Beach, 1994), we used p15 as a reporter gene to assess the induction of senescence. Knockdown of *ACVR1B* impaired the induction of p15 whereas knockdown of *TGFR1B* had no effect (Sup Fig 6d). Thus, impairment of 60S biogenesis through LSG1 knockdown elicits a restricted SASP centred around TGFβ/activin signaling.

**Fig 5.**
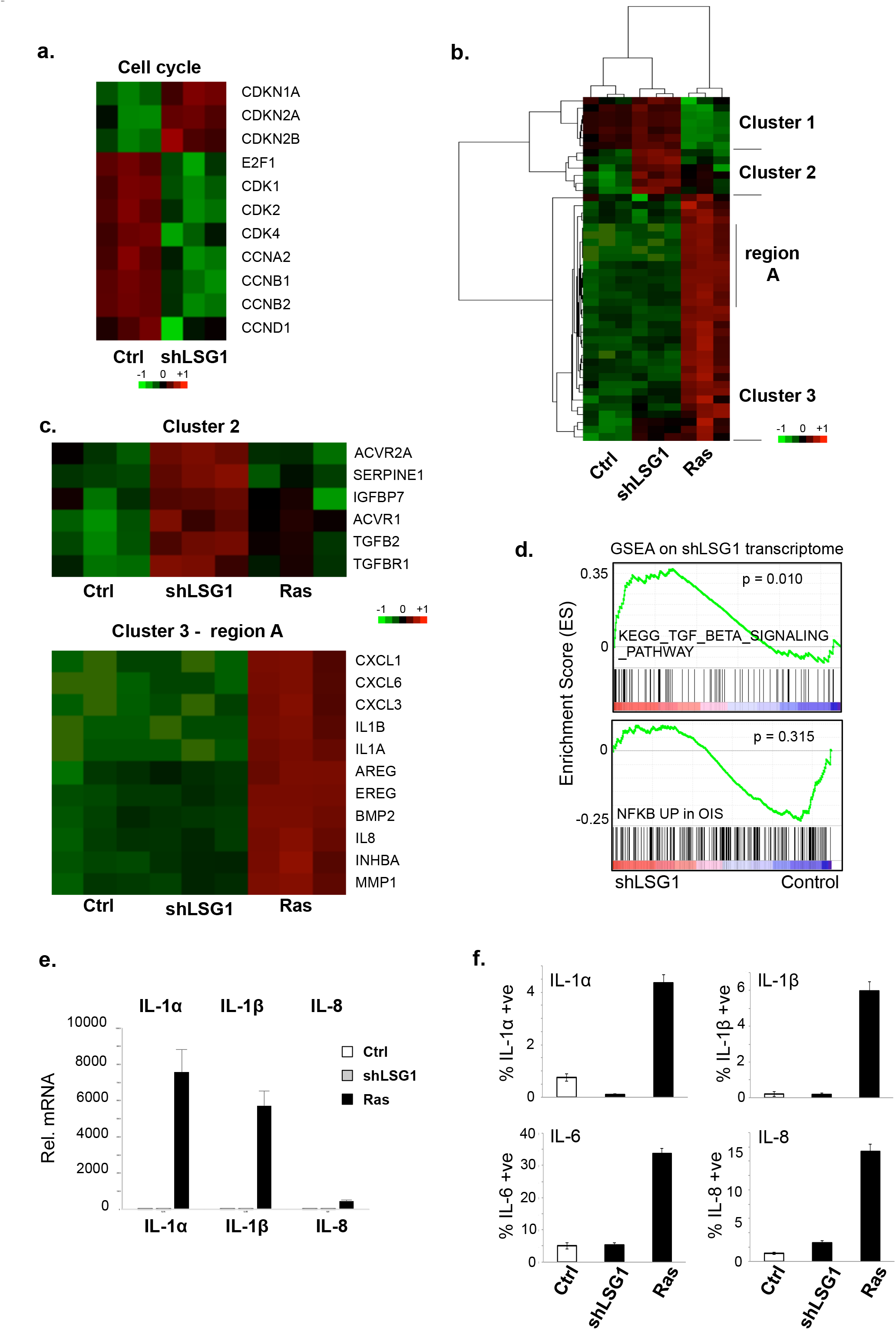
Transcriptomic analysis reveals a robust senescent transcriptional response with a restricted SASP upon LSG1 knockdown. a. Regulation of anti-proliferative and proliferative cell cycle-related transcripts by shLSG1 in MRC5 cells.
b. Clustering of transcript levels of SASP factors.
c. Cluster 2 contains a set of mRNAs that are specific for shLSG1 (versus K-RAS^V12^) that includes TGFB2 and related genes. Cluster 3, region A is OIS-specific and is comprised of NF-κB-driven canonical SASP genes.
d. GSEA of the transcriptome of MRC5 cells transduced with shLSG1 compared to control showing significant enrichment for the TGFB signalling pathway (KEGG pathway), and no significant enrichment for the OIS-associated NFKB signature (Chien et al., 2011)
e. qPCR analysis of the above cells for the quantitation of IL-1α, IL-1β and IL-8 transcript levels.
f. High content imaging analysis of the SASP factors IL-1α, IL-1β, IL-6 and IL-8. K-RAS^V12^ retroviral overexpression was included as a positive control. Error bars show standard deviation of 3 biological replicates.

### Cells in which 60S maturation is impaired are translationally active

Since LSG1 catalyses a key step in the maturation of the 60S ribosomal subunit, a possible mechanism for generation of stress leading to senescence could be a lack of 60S subunits and consequent translational insufficiency. We transduced cells with vector, shLSG1 or K-RasV12, awaited the onset of senescence and then performed polysome profiling to assess the ribosomal composition of the cells. We observed no clear differences between the conditions, except perhaps for a marginal reduction in peak height for the 60S subunit upon shLSG1 transduction (Fig 6a), suggesting that senescence occurred well before impairment of 60S biogenesis could affect overall ribosomal composition. In order to assess the impact on translation more directly, we used O-propargyl puromycin (OPP) to label actively translating ribosomes and we quantified OPP incorporation by high content microscopy. Rather than causing a reduction, knockdown of LSG1 gave rise to an elevated translation rate (Fig 6b). H-Ras^G12V^ also led to increased translation, consistent with previous reports that senescence is a cellular state associated with high translational and metabolic activity (Dörr et al., 2013; Herranz et al., 2015; Laberge et al., 2015; Narita et al., 2011). We harvested the polysomal fractions from our profiling experiment and performed qRT-PCR to assess whether mRNAs involved in the senescence response were being actively translated. As expected, polysome-associated mRNAs for p16 and p21 were elevated in both of the senescent conditions whereas IL-1α, the master regulator of the SASP (Laberge et al., 2015), was associated with polysomes in the OIS sample alone (Fig 6c). Thus the senescence response to impairment of 60S ribosomal subunit maturation is not triggered by translational insufficiency.

**Fig 6.**
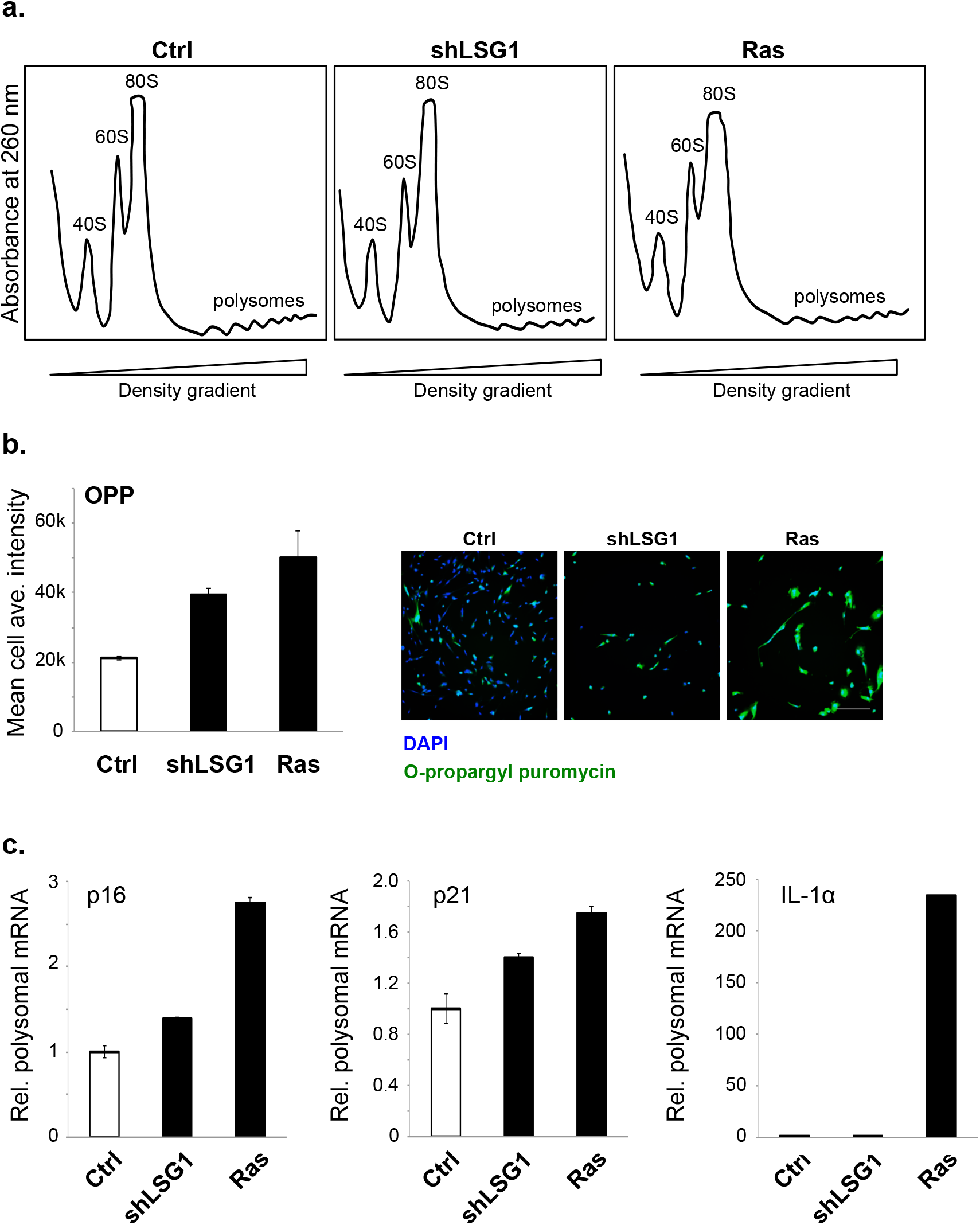
Knockdown of LSG1 does not inhibit global translation. a. Polysome profiling of MRC5 cells at senescence triggered by shLSG1 or K-RAS^V12^.
b. Analysis of translational activity using O-propargyl puromycin (OPP) and high content imaging. Quantitation of mean cell average intensity from images obtained. Representative images are provided. Scale bar: 200 μm.
c. qPCR analysis of polysome-associated transcripts for the senescence markers p16, p21 and IL-1α. Error bars show standard deviation of 3 biological replicates.

### 60S inhibition leads to disruption of ER homeostasis and morphology

In addition to the targeted analyses of transcriptomic data described above, we utilised global gene set enrichment analysis software to shed light upon the cellular response to LSG1 knockdown. This analysis revealed a striking upregulation of processes that occur at the endoplasmic reticulum (top five processes shown in Fig 7a), in particular the cholesterol biosynthesis pathway (Fig 7b). Indeed, 7 of the top 25 upregulated genes in our analysis encoded members of the cholesterol synthesis pathway (Sup Fig 7). and we confirmed upregulation of squalene epoxidase (SQLE) and hydroxymethylglutaryl-CoA synthase (HMGCS1) at the protein level (Fig 7c). The striking enrichment of the cholesterol biosynthesis and other ER-related pathways in shLSG1-induced senescence led us to look more closely at the morphology of the ER. LSG1 has been reported to predominantly localize to the ER (Reynaud et al., 2005) and its reaction partner RPL10 (also known as QM protein) has been shown to interact with ER-associated ribosomal particles (Loftus et al., 1997). Immunofluorescent staining for the ER marker calnexin revealed the expected reticular morphology of the ER in control cells, but in shLSG1 cells where NMD3 was cytoplasmic, the ER appeared highly fragmented and punctate (Fig 7d). We quantified this effect using the MiNA plugin for ImageJ (Valente,Maddalena, Robb, Moradi, & Stuart, 2017) which analyses reticularity of cellular features. Upon knockdown of LSG1, we observed a reduction in ER footprint, number of individual ER components and number of ER networks, indicating a marked disruption of ER morphology (Fig 7e). Thus, knockdown of LSG1 leads to disruption of ER homeostasis and morphology.

**Fig 7.**
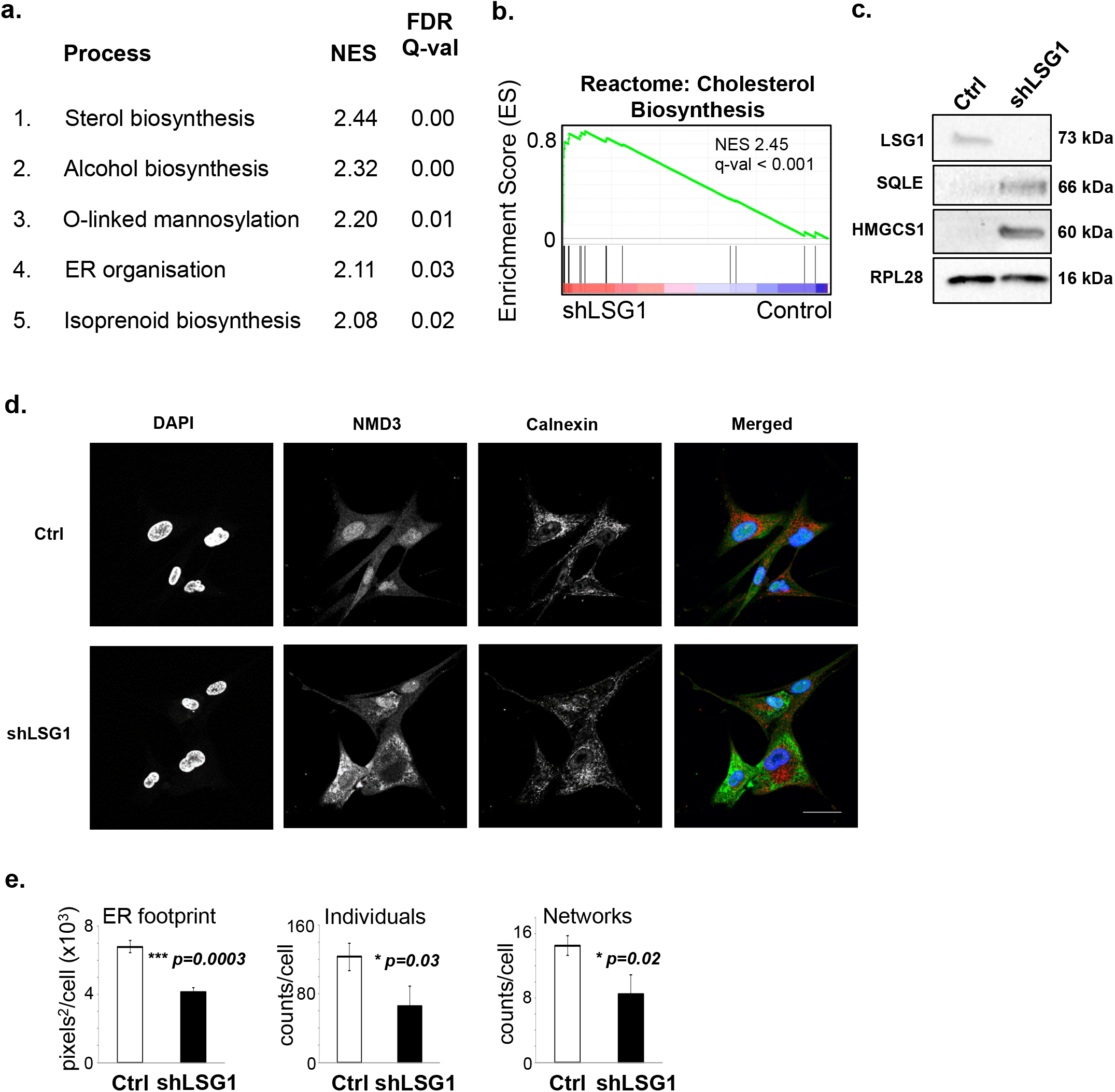
Knockdown of LSG1 leads to upregulation of cholesterol biosynthesis pathways and homeostatic alterations in the ER apparatus. a. Gene Set Enrichment Analysis (GSEA), ranked by normalised enrichment score (NES), revealed the top 5 upregulated biological processes as a result of LSG1 knockdown. The false discovery rate (FDR) yields the Q-value for statistical significance.
b. GSEA diagram of the cholesterol biosynthesis signature upon LSG1 knockdown as described in (a).
c. c. Western blot for LSG1, RPL28 and the cholesterol biosynthesis enzymes SQLE and HMGCS1 in shLSG1-transduced MRC5 cells.
d. Immunofluorescence staining for calnexin in MRC5 cells transduced with control and with shLSG1, imaged by confocal microscopy. Scale bar: 50μm.
e. FIJI-based analysis of the ER skeleton in the cells above, using the MiNA plugin (Valente,Maddalena, Robb, Moradi, & Stuart, 2017). Error bars denote SEM of three biological replicates. p-values were calculated using Student’s t-test.

### The cholesterol biosynthetic and ER transcriptomic programmes are common to the senescence induced by inhibition of 60S maturation and OIS

OIS is driven by multiple cellular stress responses, including replication stress and DNA damage, metabolic and oxidative stresses (reviewed in (Kuilman, Michaloglou, Mooi, & Peeper, 2010)). We wished to ascertain whether we could detect signals of a stress response in our transcriptomic data that were conserved between shLSG1-induced and oncogene-induced senescence (OIS). We therefore compared the transcriptomes of cells that underwent senescence due to knockdown of LSG1 or OIS induced by H-RasV12 to find genes that were upregulated in both cases. We found 125 genes upregulated in common between the two senescent programmes (Fig 8a) and we subjected these genes to gene ontology analysis. Strikingly, by far the most significant signature that emerged (Fig 8b) was cholesterol biosynthesis (p-value = 1.13 x 10^-9^), followed by ER compartment (p-value = 8.73 x 10^-4^). Since the gene sets for ER include most of the genes involved in cholesterol biosynthesis, the predominant shared component of the senescent transcriptomic response is an induction of cholesterol biosynthesis. We found that almost every gene in the cholesterol biosynthesis pathway was upregulated in both forms of senescence (Fig 8c), suggesting that the pathway may be of functional importance in the senescence response. We therefore undertook a restricted cholesterol biosynthetic siRNA screen for bypass of OIS, which we defined as an increase of 30% in BrdU incorporation compared to the senescent state. Several siRNAs from the pathway bypassed OIS (Sup Fig 8a) and the three strongest candidates from the screen (*MSMO1*, *MVD* and *DHCR7*) showed robust and significant bypass of OIS (Fig 8d, Sup Fig 7b). Thus activation of the cholesterol biosynthesis pathway is a tumour suppressive response that contributes to senescence induced by perturbation of 60S maturation and oncogenic Ras.

**Fig 8.**
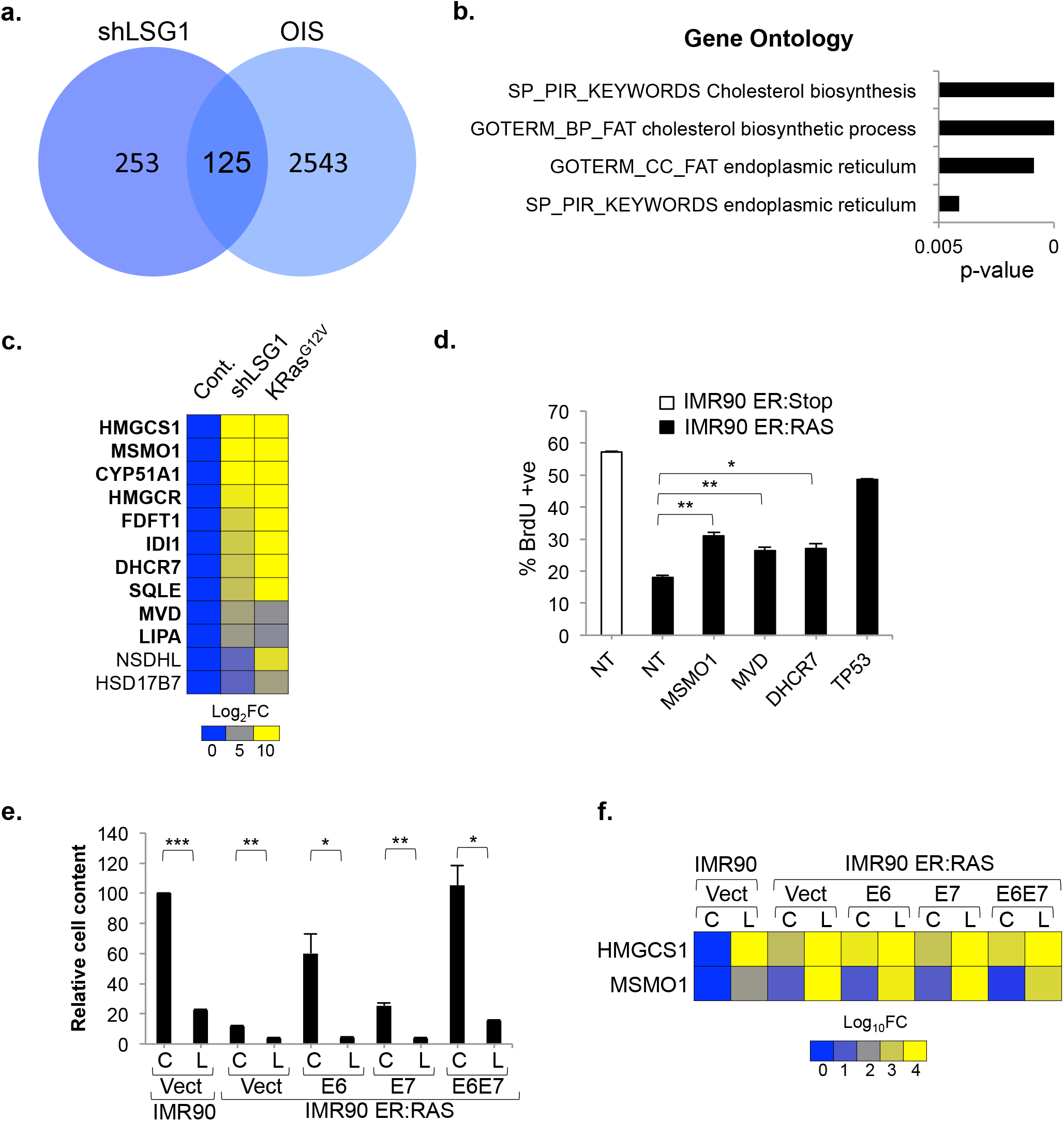
LSG1 targeting restores the cholesterol-ER senescent program in H-RAS^V12^-expressing cells that have bypassed senescence. a. Venn diagram representing the number of genes commonly induced between shLSG1 knockdown induced senescence and OIS in MRC5 cells by Ampliseq transcriptome analysis.
b. Bar graph representing the p-value after functional annotation analysis of the most significant GO terms enriched in the 125 genes induced by shLSGI and oncogenic RAS in MRC5 cells as in a. Analysis was performed using the DAVID web resource (https://david.ncifcrf.gov).
c. Heat map representing mRNA fold change (Log2 scale) in the Ampliseq expression profile of cholesterol biosynthesis genes in shLSG1 and RAS transduced MRC5 cells. Each sample represents the mean of 3 experimental replicates. Bold character genes represent significant changes in expression in both conditions.
d. . BrdU proliferation assay of IMR90 ER:RAS or ER:Stop control cells 5 days after 4 hydroxytamoxifen (4OHT) treatment and siRNA smartpool transfection for the cholesterol biosynthesis genes MSMO1, MVD, DHCR7 and TP53 (as a positive control). Non-targeting (NT) siRNA smartpool was use as a negative control. Bars represent the mean of 3 experimental replicates. Error bars represent the SEM.
e. Proliferation assay showing relative cell content of cells transduced with shLSG1 or control lentiviral vectors in cells bypassing OIS. Bypass of OIS was achieved with retrovirus expressing HPV proteins E6, E7, E6E7, or neomycin control. Cells were seeded at low density, cultured for 14 days and stained with crystal violet (CV) as indicated. Bars represent the mean quantification of CV staining of three independent experiments. Error bars represent the SEM.
f. Heat map showing HMGCS1 and MSMO mRNA fold change (Log_10_ scale) by RT-qPCR from cells treated as in (e) above.

### shLSG1 amplifies the cholesterol biosynthesis signature and induces senescence in cells that have bypassed OIS

A critical step in the transformation of cells expressing oncogenic Ras is the bypass of OIS through disruption of the p53 or RB pathways (Serrano et al., 1997). One or both of these two canonical tumour suppressive pathways is inactivated in most cancers and thus, for a prosenescent cancer therapy to be effective, it should be able to elicit tumour suppression independently of these two pathways. We wished to assess whether the induction of the cholesterol biosynthesis programme by inhibition of 60S maturation might provide such a tumour suppressive response. We therefore generated pre-transformed cells through a combination of overexpression of H-RasV12 and the human papilloma virus oncoproteins E6 (which inactivates p53), E7 (which inactivates RB) or an E6-E7 fusion and then performed knockdown of LSG1. We observed that knockdown of LSG1 reduced cell content in the absence or presence of oncogenic Ras and that E6, E7 and E6-E7 bypassed Ras-induced growth arrest (Fig 8e, Sup Fig 8c), as expected. In the conditions where OIS was bypassed, shLSG1 elicited a marked growth arrest, even in the E6-E7 line where both p53 and RB pathways are defective (Fig 8e, Sup Fig 8c). This growth arrest was accompanied by induction of the cholesterol biosynthesis pathway (shown for *HMGCS1* and *MSMO1* in Fig 8f) and acidic β-galactosidase staining indicated that the reduced cell number was due to a senescence response (Sup Fig 8d). Taken together, these data reveal that inhibition of 60S maturation restores a tumour suppressive senescence response even in cells that have bypassed OIS.

## Discussion

Here we show that inhibition of 60S maturation leads to a robust induction of cellular senescence through perturbation of ER homeostasis and that this can elicit tumour suppression even in cells with bypass of OIS. The impairment of 60S ribosomal subunit maturation upon knockdown of LSG1 was verified by relocalisation of NMD3 to the cytoplasm in analogous fashion to the response to disruption of *Lsg1* in *S. cerevisiae* (Hedges et al., 2005). However, rather than causing accumulation of pre-60S subunits and decreased polysomes as in *S. cerevisiae*, it resulted in an increase in translation accompanied by normal ribosome content, consistent with previous reports of senescence as a highly metabolically active process requiring elevated rates of translation (Dörr et al., 2013; Narita et al., 2011). Similarly, deletion of *Sbds* in normal mouse pancreas was recently shown to elicit a senescent response without perturbation of global ribosome content (Tourlakis et al., 2015), whilst the equivalent perturbation in *S. cerevisiae* promotes impairment of the polysome profile (Menne et al., 2007). Taken together, these reports suggest that an important function of the senescence response may be to halt cellular proliferation prior to the onset of a translational defect, thereby protecting cellular translational capacity in response to perturbations of ribosome biogenesis. *S. cerevisiae* lacks the ability to mount complex stress responses such as senescence and therefore these 60S defects result in catastrophic reduction of ribosome content.

Senescence is a pleiotropic response to many cellular stresses and, although many of the effector pathways (e.g. cell cycle arrest, the SASP etc.) are well characterised, the precise molecular mechanisms that trigger senescence remain obscure in most cases. Transcriptomic analyses can shed light upon molecular mechanisms of cellular stresses because discrete effector pathways often reveal the nature of the initial stress, for example the induction of NRF-2 gene targets in response to oxidative stresses (reviewed in (Nguyen, Nioi, & Pickett, 2009)) and HIF-1 gene targets upon hypoxia (reviewed in (Kaluz, Kaluzová, & Stanbridge, 2008)). Our transcriptomic analyses gave a clear indication of stress arising at the ER and our further analyses revealed disruption of ER morphology upon loss of LSG1. The origin of a cellular stress response at the ER is consistent with previous reports of LSG1 and RPL10 localisation and function at the ER (Loftus et al., 1997; Reynaud et al., 2005). It is unlikely that the LSG1/RPL10-mediated removal of NMD3 only occurs at the ER and we favour a model whereby this reaction occurs throughout the cytosol and at the ER, but the stress response arises due to perturbation of the latter. At this time, it is unclear why there is such a specific activation of the cholesterol biosynthesis pathway by shLSG1 and why this signature is also so prevalent in OIS and other forms of senescence. A recent study identified accumulation of the ribosomal 40S subunit protein RPS14 as a mechanism contributing to senescence in response to multiple stimuli (Lessard et al., 2018). Unlike our p53-dependent response, the response to RPS14 was Rb-dependent, indicating that it is mechanistically distinct. It therefore seems that cells use multiple mechanisms to surveil ribosome biogenesis and that senescence is the outcome when defects are detected.

The inhibition of 60S ribosomal subunit maturation gives rise to a robust senescence response that is comparable with the OIS induced by K-Ras^V12^ in all aspects that we examined except for the SASP. The SASP elicited by deregulation of Ras in fibroblasts is a cocktail of proinflammatory cytokines and chemokines resembling those produced during an immune response to infection. On the other hand, the restricted SASP activated upon inhibition of 60S maturation primarily involves components of the TGFβ signalling pathway. In terms of a potential cancer therapy, the absence of a strongly pro-inflammatory SASP is likely to be a considerable advantage, since pro-inflammatory signaling, through IL-6 in particular, has been linked to tumour progression and metastasis (He et al., 2013; Kim et al., 2009).

A therapeutic concept that is supported by our data is that inhibition of ribosome biogenesis could be an effective cancer therapy and our induction of tumour suppression in cells with defective p53 and RB pathways is particularly encouraging. GTPases have not previously been strong candidates for inhibition through small molecules, although the translational GTPase eEF2, a homologue of EFL1, has been well validated as an inhibitory target since the naturally occurring inhibitors sordarin, diphtheria toxin and exotoxin A all target this enzyme. Inhibition of eEF2 is toxic to mammalian cells due to inhibition of translation, but here we demonstrate that inhibition of LSG1, and possibly EFL1 by extension, may provide an effective prosenescent cancer therapy with lesser side effects since translation remains unimpaired. Recently, an important advance in the field of GTPase inhibition was reported with the identification of a non-nucleotide active site inhibitor of the small GTPase Rab7 that can act as a scaffold for derivatisation to produce inhibitors of other GTPases (Agola et al., 2012; Hong et al., 2015). Accordingly, the translational GTPases of 60s ribosomal subunit biogenesis may be amenable to development of inhibitors. In conclusion, this study suggests that the GTPase LSG1 has high potential as a candidate target for pro-senescent cancer therapy in cases where tumour suppressive senescence is bypassed due to p53 and/or RB deficiency.

## Methods

### Cell culture, viral transduction and siRNA transfection

MRC5 and IMR90 early passage primary human fibroblasts were purchased from the Culture Collection at Public Health England. These cells and HEK293ET (used for viral production – a kind gift of Felix Randow at the MRC Laboratory of Molecular Biology, Cambridge) were propagated in DMEM with added 10% FCS and 5% Pen/Strep. Lentiviral production for shRNAs was carried out by transfection of HEK293ET cells with a packaging vector vector (psPAX2), VSV-G envelope (pMD2.G) and viral transfer vector (listed below). For retroviral transduction, pGag-Pol was used in place of psPAX2. Typically 10 μg of each vector was combined with 80 μl of polyethyleneimine (PEI - 1 μg/μl) in a 500ul volume (the remainder being DMEM). This was then added to a 75 cm^2^ flask of cells containing 10 ml of DMEM/10% FCS and incubated overnight. Medium was exchanged the following day for DMEM/10% FCS and left for a further 24 hours at which point the viral supernatant was harvested for infection. Viral supernatant was diluted (typically 3:10) with DMEM/10% FCS and mixed with polybrene (hexadimethrene bromide) at a final concentration of 5 μg/ml. This was filtered through a sterile 0.45 μm filter and used to replace the medium on recipient cells. 48 hours after infection, antibiotic was added for selection, typically puromycin at 1 μg/ml or blasticidin at 5 μg/ml, and cells were selected until uninfected control cells had died. Timepoints referred to are days post-infection (not selection). For siRNA transfections, plated cells were transfected with a mix of medium containing SmartPool siRNA (Dharmacon) at 50nM and 3.5% Hiperfect transfection reagent (Qiagen).

### Viral transfer vectors

Knockdown of 60S maturation factors using lentivirus was carried out using pLKO1 or Tet-LKO-puro containing oligos as follows:

Ctrl: CCGGTCCGCAGGTATGCACGCGTG

LSG1: CCGGTGGGCTACCCTAATGTTGGTACTCGAGTACCAACATTAGGGTAGCCCATTTTTG

SBDS(a): CCGGAAGCTTGGATGATGTTCCTGACTCGAGTCAGGAACATCATCCAAGCTTTTTTTG SBDS(b): CCGGCTGCTTCCGAGAAATTGATGACTCGAGTCATCAATTTCTCGGAAGCAGTTTTTG. E6, E7

and E6E7 constructs in pLXSN have been previously described (Juan CAcosta et al., 2008). Dominant negative p53 (Genbank KF766124) and KRasV12 were expressed in the retroviral vector pM6P-Blast (a kind gift of Felix Randow, MRC Laboratory of Molecular Biology).

### Western Blotting and Antibodies

Blots were performed according to standard protocols (with overnight incubation of primary antibody at 4°C). Antibodies used were raised against: LSG1 (Proteintech 17750), EFL1 (24729), SBDS (Abcam ab128946), RPL28 (Proteintech 16649), BrdU (Pharmingen 558599), p53 (Santa Cruz sc-126), p16 (Santa Cruz sc-56330), p21 (Sigma p1484), pST/Q (Cell Signaling 2851), IL-1α (R&D MAB200), IL-1β (R&D MAB201), IL-6 (R&D AF206NA), IL-8 (R&D MAB208), Ki67 (Invitrogen 180191Z), E-Cadherin (BD Biosciences 612130).

### High Content Microscopy

High content microscopy was performed as previously described (Hari & Acosta, 2017). Where included, 50 mM BrdU was incubated with cells for 16 hours prior to fixation. Briefly, cells in 96-well plates were fixed with 10% formalin for 10 mins and then blocked with blocking solution (1% BSA/0.2% fish gelatin in PBS). Primary antibody was then added and cells were incubated for 1 hr at RT. Anti-BrdU solution was supplemented with 0.5U/μl DNAse (Sigma D4527) and 1 mM MgCl_2_. Following incubation with fluorescent secondary antibodies for 1 hr and 1 μg/ml DAPI for 30 min, plates were loaded onto an ImageXpress Micro High Content Imaging System (Molecular Devices) and fluorescent images were acquired. Results were analysed using MetaXpress software (Molecular Devices).

### Cytochemical staining for SA-β-galactosidase

Cell fixation was performed in 0.5% glutaraldehyde/PBS for 15 minutes at RT. After washes in 1 mM MgCl_2_/PBS, pH 6, the cells were incubated in staining solution [2 mg/ml 5-bromo-4-chloro-3-indolyl-β-D-galactopyranoside (Sigma, B4252), 1.64 mg/ml K_3_Fe(CN)_6_, 2.1 mg/ml K_4_Fe(CN)_6_.3H_2_O in 1mM MgCl_2_ 1, pH 6] at 37 °C, for 24 hours. The production of a blue precipitate within the cytoplasm, as observed under an inverted microscope, determined the lysosomal SA-β-gal activity (Dimri et al., 1995).

### O-Propargyl Puromycin (OPP) assay

For the OPP assay we used the Click-iT Plus OPP Alexa Fluor 488 kit (ThermoFisher) and followed the manufacturer’s protocol. The cells were labeled with OPP at a final working solution of 20μM for 30 min. For OPP detection standard immunofluorescence procedures were followed and the cells were acquired using a high content screening automated microscope.

### Transcriptomic analysis

RNA was harvested from cells using an RNEasy/QIAshredder (Qiagen) protocol following the manufacturer’s instructions. Reverse transcription of DNA was carried out using QScript enzyme (Quanta) and RNA was submitted to the Genome analysis core at the Wellcome Trust Clinical Research Facility (Western General Hospital) for AmpliSeq library preparation and IonTorrent sequencing. Analysis of the data was performed using the Babelomics-5 application (http://babelomics.bioinfo.cipf.es). Hierarchical clustering analysis was performed using Cluster 3 software (Stanford University) and visualization was performed using TreeView 3.0 software (Princeton University).

### Polysome profiles

Cells were lysed in detergent lysis buffer A (10 mM Tris-HCl at pH 7.4, 10 mMNaCl, 1.5 mM MgCl2, 0.5% [v/v] Triton X-100, 0.5% [w/v] deoxycholate, 1% [v/v] Tween 20, 100 mg/mL cycloheximide) with complete EDTA-free protease inhibitors (Roche) and 0.5

U/mL RNase inhibitor (Promega) and incubated for 10 min on ice. Lysates were cleared in a microfuge. Equal amounts (typically 10–20 A254 U) were applied to a 10%–50% (w/v) sucrose gradient in 11 mL of buffer B (10 mM Tris-HCl at pH 7.4, 75 mM KCl, 1.5 mM MgCl2) and centrifuged (Beckmann SW41 rotor) at 41,000 rpm for 80 min at 4°C. Samples were unloaded using a Brandel gradient fractionator, the polysome profiles were detected using a UV monitor (Gilson) at A254, and fractions were collected. For analysis of polysome-associated mRNAs, polysomal fractions were pooled and RNA was purified using the RNEasy mini kit (Qiagen). RNA was reverse transcribed with QScript enzyme (Quanta) according to the manufacturer’s protocol and the cDNA was used as a template for PCR. Primer oligos used were:

p16: CGGTCGGAGGCCGATCCAG / GCGCCGTGGAGCAGCAGCAGCT

p21: CCTGTCACTGTCTTGTACCCT / GCGTTTGGAGTGGTAGAAATCT

IL1α: AGTGCTGCTGAAGGAGATGCCTGA / CCCCTGCCAAGCACACCCAGTA

β-actin: CATGTACGTTGCTATCCAGGC / CTCCTTAATGTCACGCACGAT

### CRISPR and focus formation assays

CRISPR-mediated disruption of the p53 gene was carried out using the vector lentiCrisprV2 (Addgene #52961) and sgRNAs for non-target (NT - ACGGAGGCTAAGCGTCGCAA) and p53 (GAGCGCTGCTCAGATAGCGA). Targeting was assessed by PCR and Surveyor assay following manufacturer’s instructions (Integrated DNA Technologies) using the following oligos:

CTAGTGGGTTGCAGGAGGTGCTTA / CAGAGACCCCAGTTGCAAACCAG

Cells were transduced with lentivirus for shLSG1 or retrovirus for K-RasV12 and seeded at low density (50,000 cells per plate). At the endpoint, plates were stained with 0.15% crystal violet and imaged for colony formation. For quantitation, crystal violet was extracted using 1M acetic acid and relatively quantified through measurement of absorbance at a wavelength of 595 nm.

## Acknowledgements

Funding for this work was from Cancer Research UK (Intermediate fellowship to JCA). We thank Dasa Longman, Maria Christophorou, Noor Gammoh and Alex von Kriegsheim for helpful discussions.

---

**Sup Fig 1.**
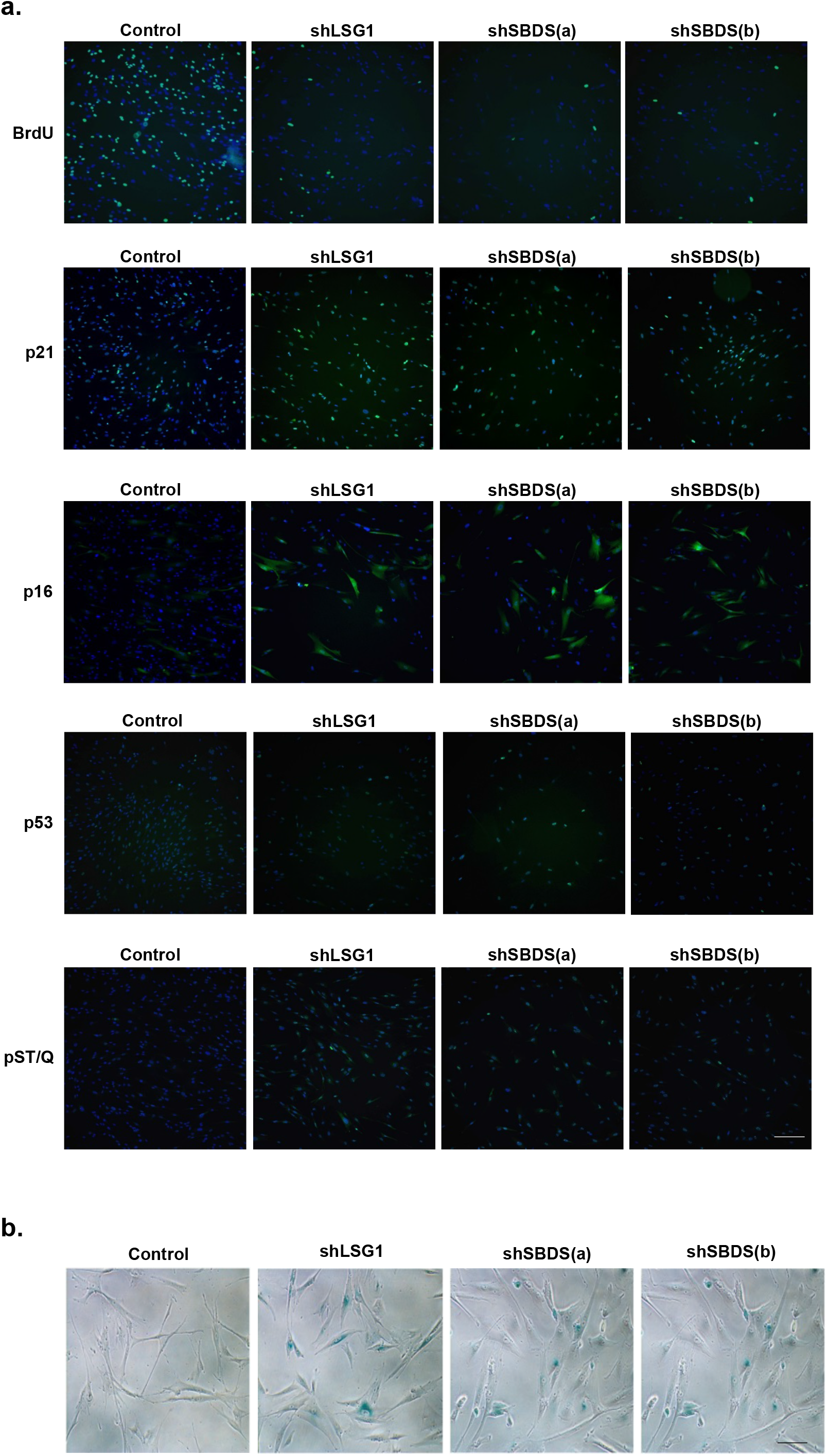
Senescence induced by shLSG1 and shSBDS is accompanied by expression of several senescence markers and high β-galactosidase activity. a. Representative images from high content imaging analysis of BrdU incorporation and immunostaining for p53, p16, p21 and the DNA damage marker pS/TQ in MRC5 cells with knockdown of LSG1 and SBDS. Scale bar: 250 μm.
b. Images from Senescence-Associated β-galactosidase assay. Scale bar: 100 μm.

**Sup Fig 2.**
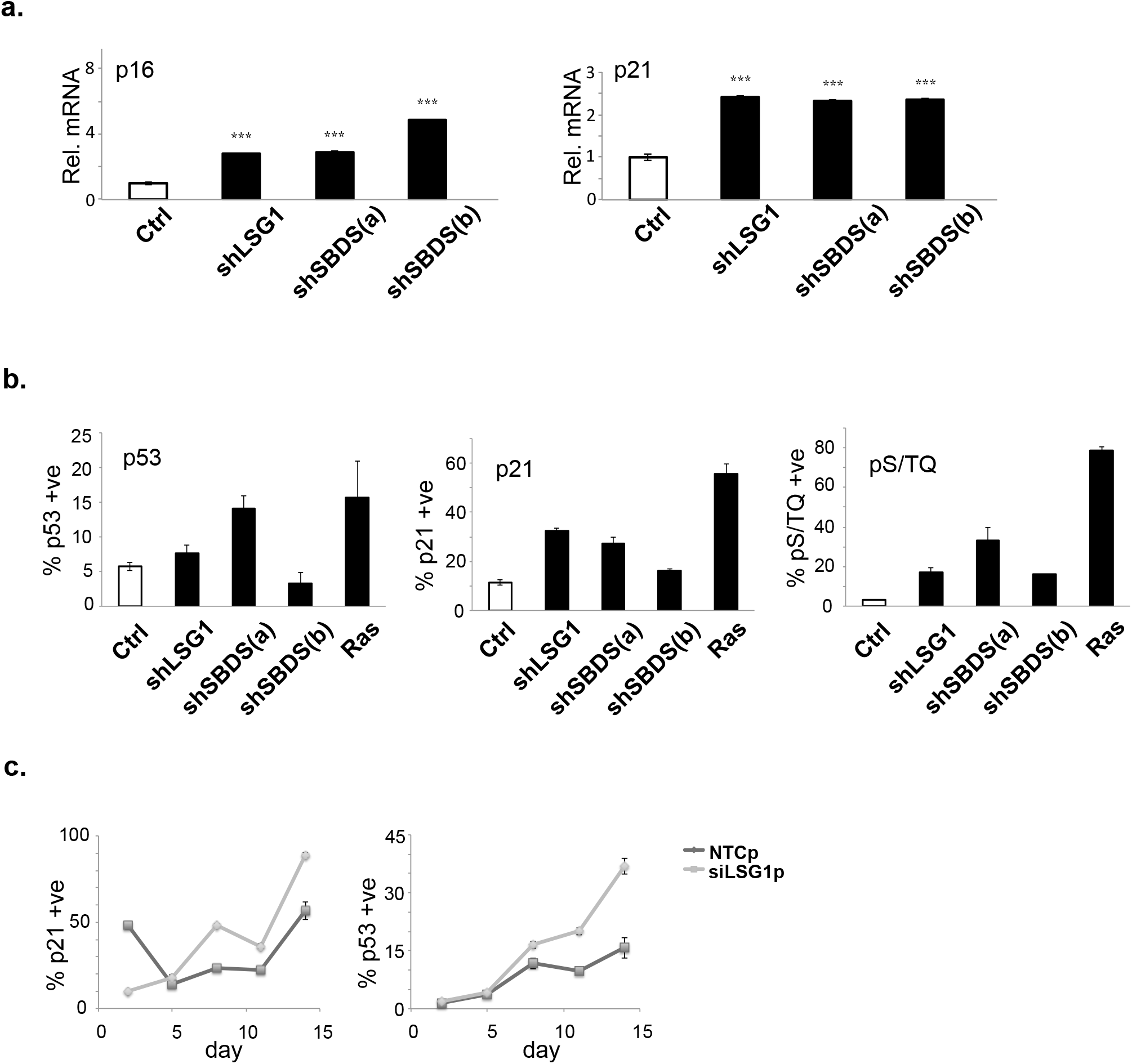
Increased mRNA and protein levels of p16, p21 and p53 in shLSG1-and shSBDS-induced senescence. a. qPCR analysis of MRC5 cells with knockdown of LSG1 and SBDS for the quantification of p16 and p21 transcript levels. Error bars show the standard deviation of 3 biological replicates.
b. High content imaging analysis of p53, p21 and pST/Q expression levels, upon immunostaining. K-RAS^V12^ retroviral overexpression was included as a positive control. Error bars show the standard deviation of 3 biological replicates.
c. Timecourse experiment (timepoints: d2, d5, d8, d11, d14) using a siRNA SMARTpool for LSG1. p21 and p53 expression levels were monitored over the course of the 14-day period, using high content microscopy. Error bars show the standard deviation of 3 biological replicates.

**Sup Fig 3.**
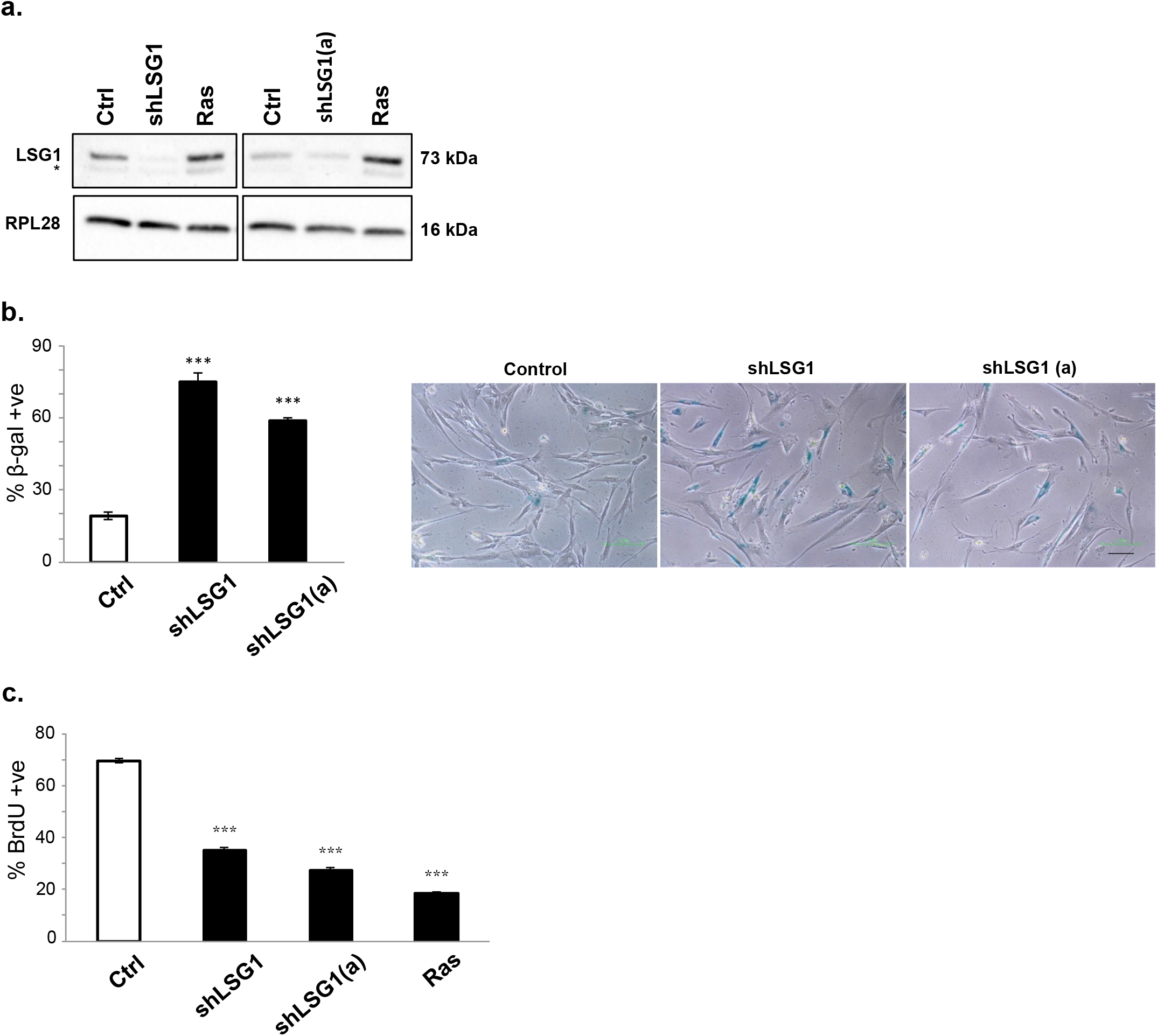
Additional shRNA against LSG1 induces senescence. a. Western blot indicating LSG1 knockdown in MRC5 cells induced by the hairpins shLSG1 and shLSG1(a). RPL28 was used as a reference protein. (The asterisk denotes a non-specific band in the LSG1 blot). K-RAS^V12^ cells were used as a positive control for senescence.
b. The Senescence-Associated β-galactosidase assay was performed 7 days post-transduction. Representative images are provided. Scale bar: 80 μm.
c. BrdU incorporation as measured by high content imaging in cells transduced as in (a) above. Error bars show the standard deviation of 3 biological replicates.

**Sup Fig 4.**
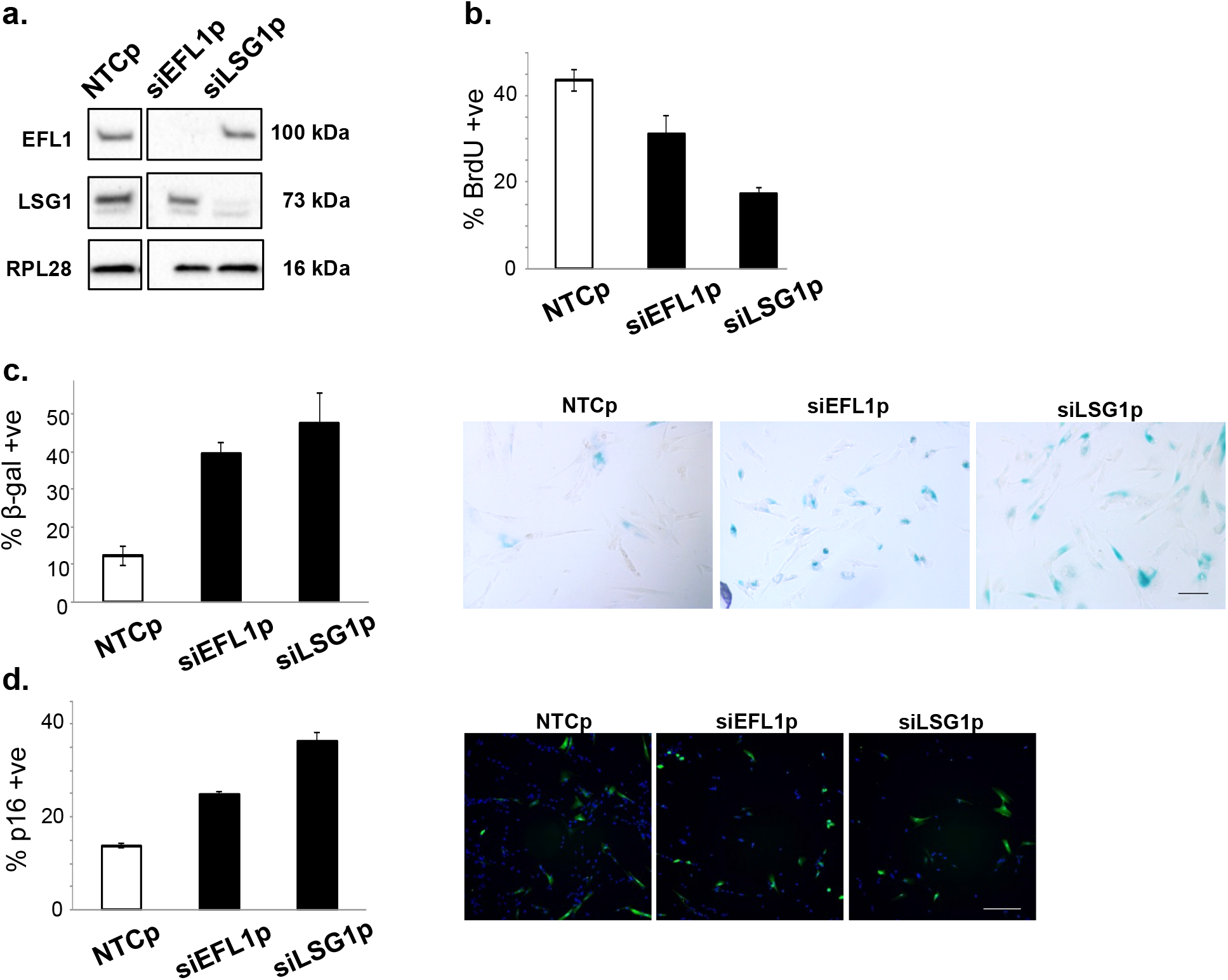
siRNA inhibition of EFL1 and LSG1 causes senescence. a. Western blot showing knockdown of EFL1 and LSG1 in MRC5 cells upon transfection with SMARTpool siRNAs.
b. High content imaging analysis of BrdU incorporation in the cells described in (a).
c. Quantitation and representative images of Senescence-associated β-galactosidase staining in the cells described in (a). Scale bar: 100 μm.
d. p16 staining in the above cells, followed by high content imaging. Representative images are shown. Scale bar: 200 μm. Error bars show the standard deviation of 3 biological replicates.

**Sup Fig 5.**
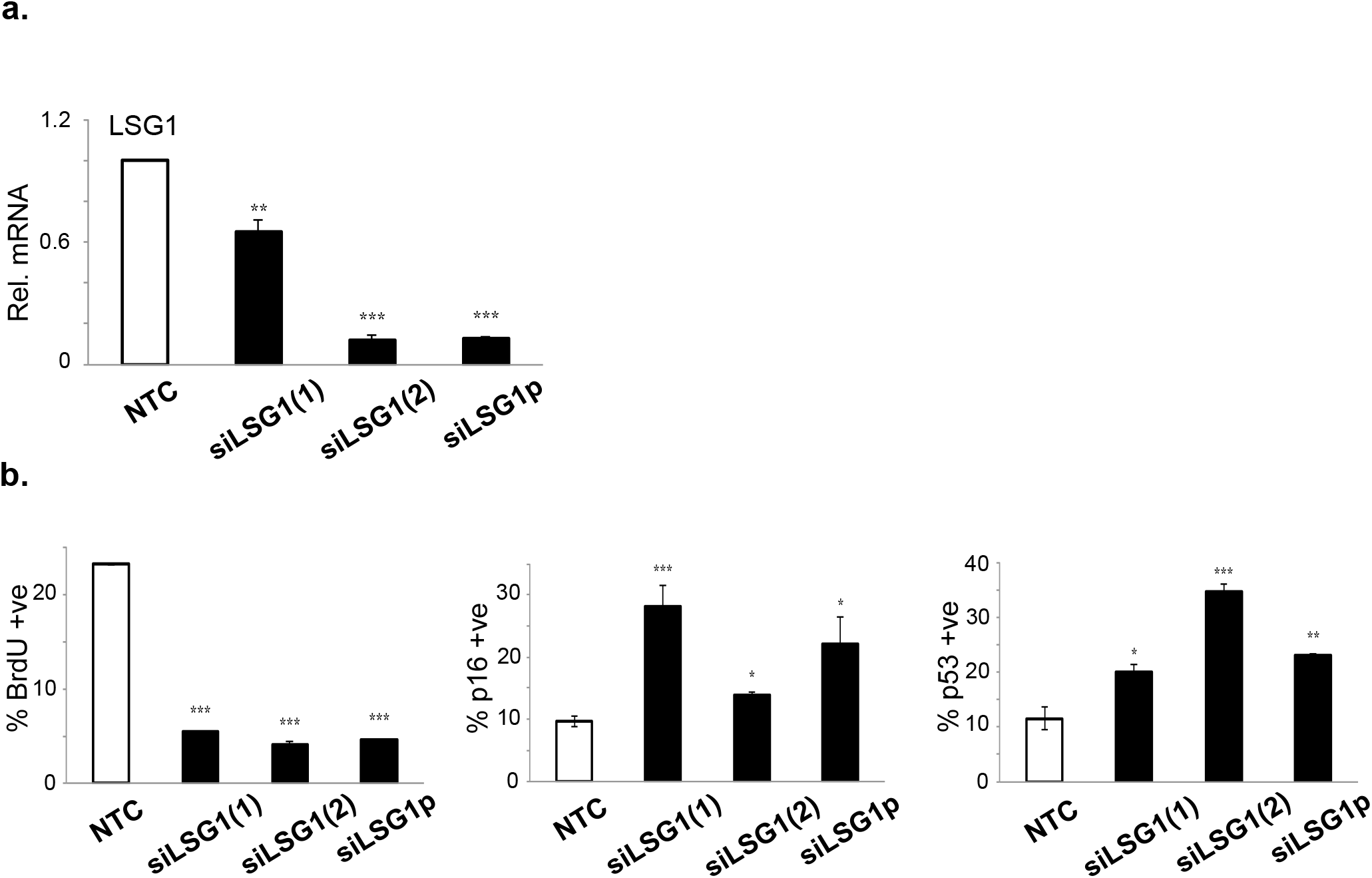
Deconvoluted siRNA inhibition of LSG1 causes senescence. a. qPCR analysis of LSG1 mRNA expression upon single transfection of component siRNAs from the SMARTpool.
b. High content imaging analysis of BrdU incorporation and p16 and p53 expression in MRC5 cells with knockdown of LSG1 mRNA. Error bars show the standard deviation of 3 biological replicates.

**Sup Fig 6.**
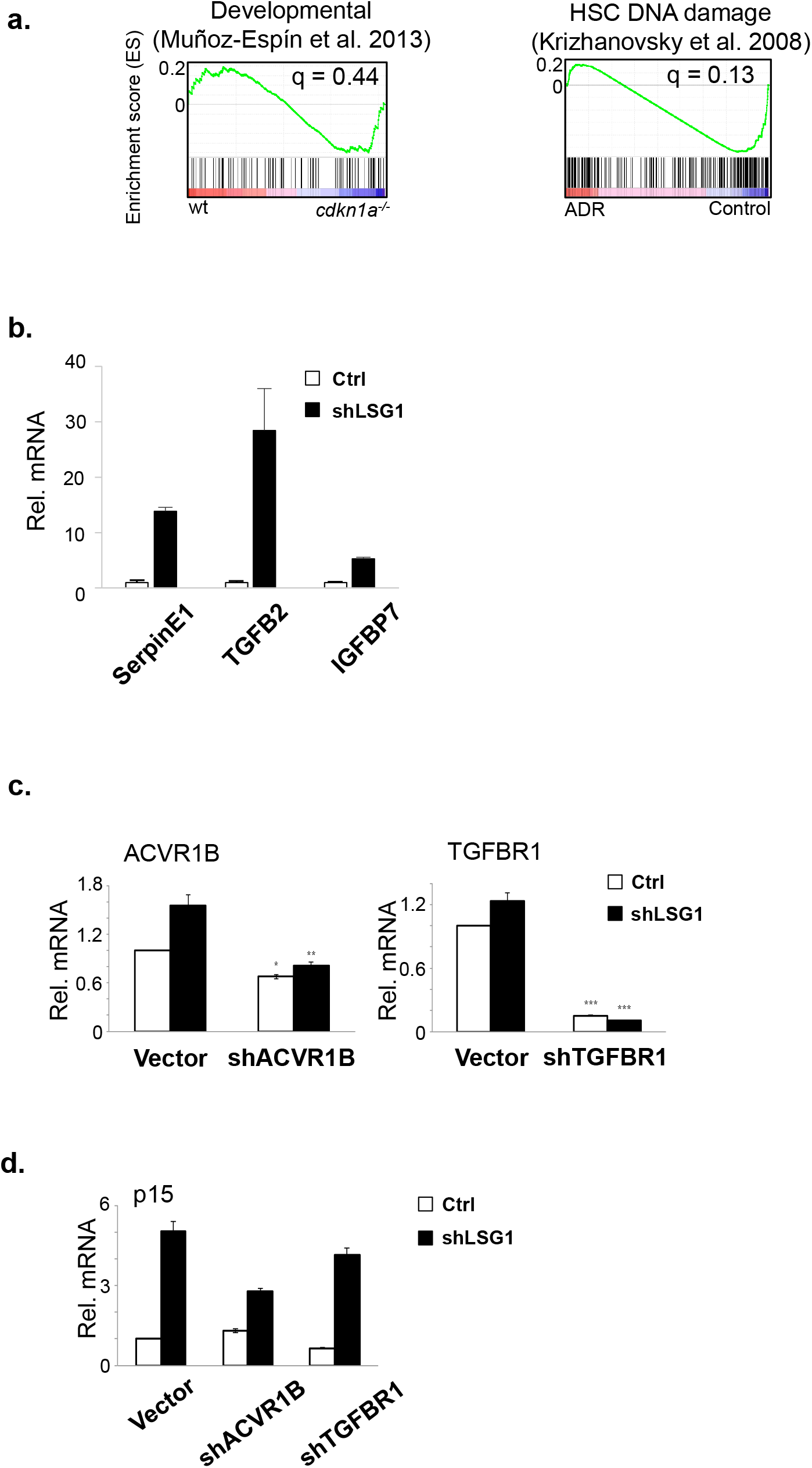
a. A signature of 253 genes derived from MRC5 cells undergoing shLSG1-induced senescence (as in Fig 4) is not significantly enriched in programmed developmental senescence (Munoz-Espin et al. 2013) and DNA damage induced senescence (Krizhanovsky et al. 2008)
b. qPCR analysis for detection of the mRNA expression levels of SerpinE1, TGFB2 and IGFBP7 in shLSG1-and vector-transduced MRC5 cells.
c. qPCR analysis for detection of the mRNA expression levels of ACVR1B, TGFBR1 in shLSG1-and vector-transduced cells infected with shACVR1B and shTGFBR1 as indicated.
d. qPCR analysis for detection of the mRNA expression levels of p15 across the several cell lines. Error bars show the standard deviation of 3 biological replicates.

**Sup Fig 7.**
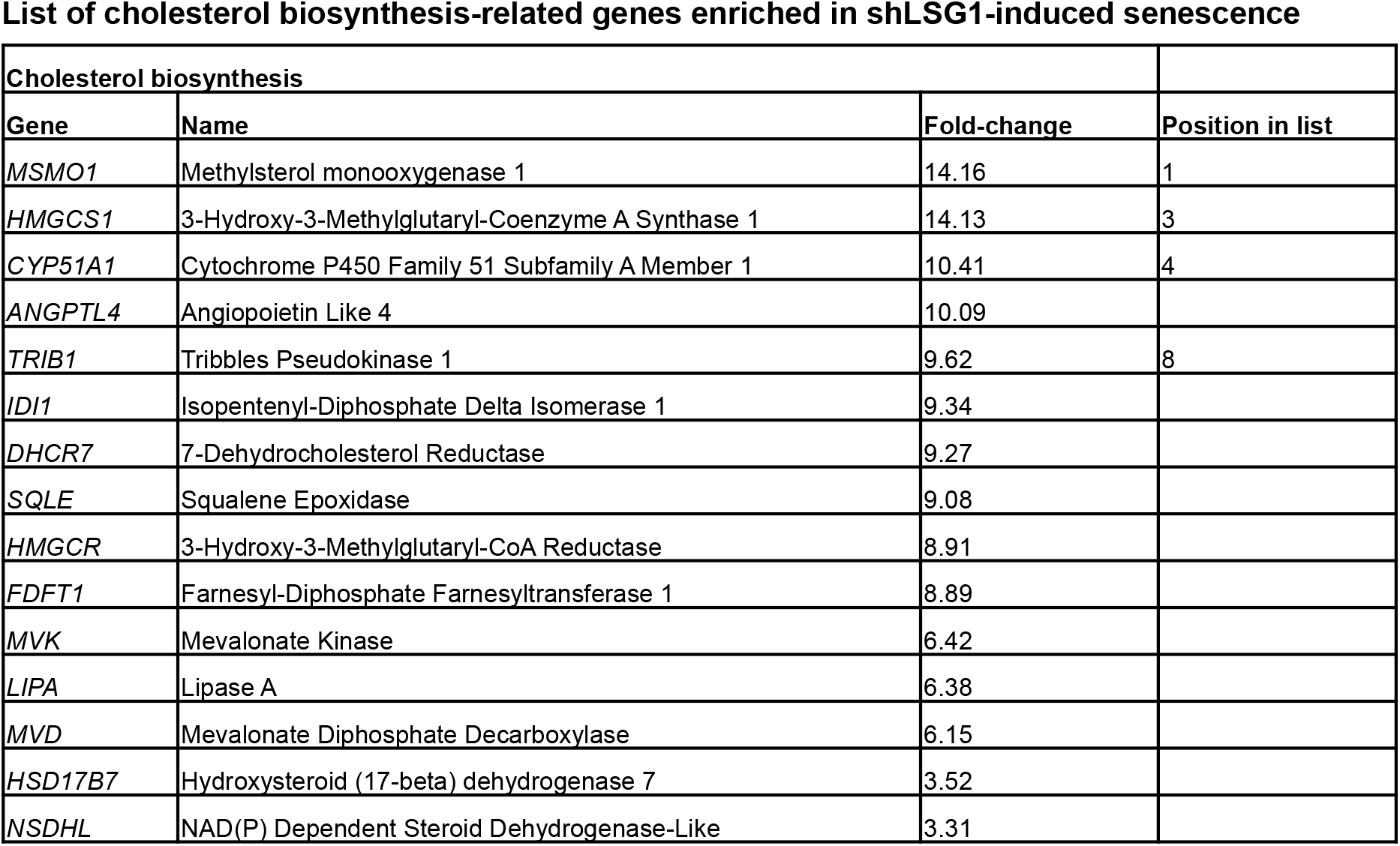
List of cholesterol biosynthesis-related genes enriched upon LSG1 downregulation. The analysis was performed using the software GSEA (Broad institute).

**Sup Fig 8.**
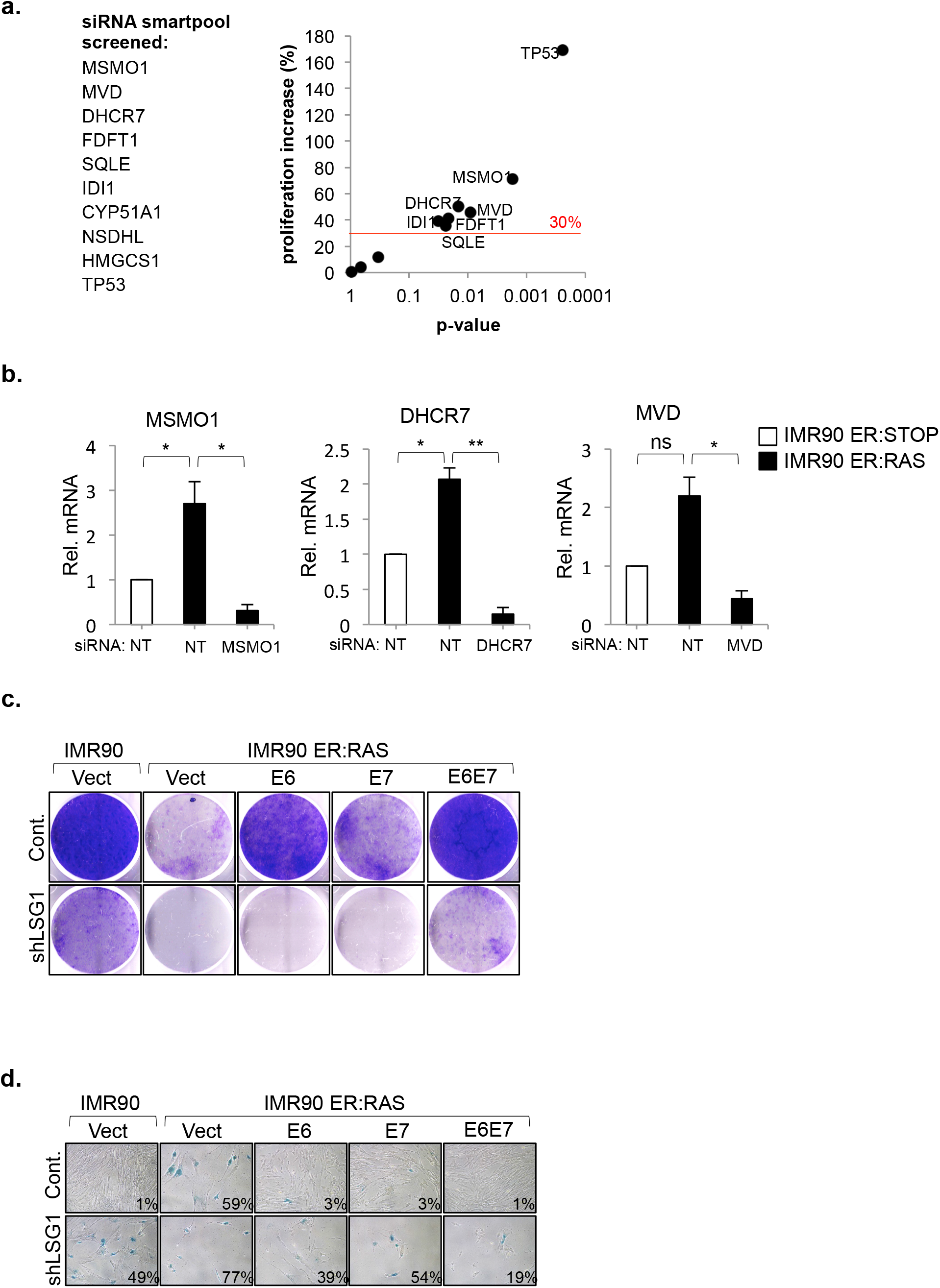
a. 9 genes regulating cholesterol biosynthesis were screened for their effect in proliferation in OIS by BrdU proliferation assay of IMR90 ER:RAS cells 5 days after 4 hydroxytamoxifen (4OHT) and siRNA smartpool transfection. Non-targeting (NT) siRNA smartpool was use as a negative control. Graph represents the increase of BrdU incorporation over the NT siRNA control versus the p-value of three independent experimental plates. The red line indicates the 30% increase in proliferation over the NT control, which was set as the threshold to be considered as a positive effect in OIS. siRNA to p53 (TP53) was used as a positive control for senescence bypass.
b. mRNA expression analysis by qRT-PCR showing knockdown of siRNA targets for the experiment described in figure 8d.
c. Representative images of tissue culture dishes stained with crystal violet from the experiment described in figure 8e.
d. Senescence associated β-Galactosidase staining from the experiment described in figure 8e.

